# Tracking the origin of island diversity: insights from divergence with the continental pool in monocots

**DOI:** 10.1101/678300

**Authors:** Simon Veron, Maud Mouchet, Philippe Grandcolas, Rafaël Govaerts, Thomas Haevermans, Roseli Pellens

## Abstract

At their early age, a large proportion of island pools were a partial sampling of mainland pools whatever islands are oceanic or fragments of the mainland. Through time, colonization, diversification, extinctions, have deeply transformed insular and continental communities and therefore the degree to which they share species. We studied the relative importance of these mechanisms by looking at the shared evolutionary history between species pools on islands and continents. Indeed, most of these mechanisms are not neutral regarding phylogeny but are linked to species evolutionary relationships. We measured the phylogenetic divergence between continental and insular monocot communities through metrics of beta Mean Pairwise Distance and beta Mean Nearest Taxon Distance. We first tested the influence of spatial and environmental distance to the continent, two main factors of divergence, but whose explanatory power in a phylogenetic context was still unclear. We showed that both dispersal and enviromental filtering were important to explain divergence, although species that could pass these filters were not phylogenetically clustered. There was however a clear distinction between oceanic and continental islands: a stable climate in the latters was key to the survival of the original biota leading to a high proportion of shared lineages between the mainland and islands. But distance to the continent was only part of the story, we investigated additionnal mechanisms of phylogenetic divergence through their relation to island features and community structure. This showed that the most divergent islands occurred in the tropics and that processes of persistence of original species, diversification in remote archipelagos, neutral colonization on easy-to-reach islands, turnover, and high speciation rates may have driven phylogenetic divergence at a world scale. This study showed how phylogenetic approaches may explain how divergence, or similarity may have arisen and provide new insights in the continental origin of plant diversity on islands.

## Introduction

One of the main questions concerning island ecology is the extent their floras and faunas resemble those from the continents assumed to be the main source of colonisers (Patiño et al. 2017, Koenig et al. 2019). At an early stage, a large proportion of island communities are a partial sampling of mainland pools (Wallace 1880 but see Ali 2017). Continental islands already harbored a full complement of species when they separated from continents. Oceanic islands were depauperate of life at birth, and their species composition results mainly from colonization from continents. Through time, large scale factors such as geographical isolation, archipelago dynamics and island age, have shaped the arrival, settlement, diversification of species in the community. In addition, community assembly in islands may be influenced by other determinants such as habitat and resource availability that modulate population sizes or inter and intra specific interactions, driving selection, survival and speciation (Kreft et al. 2008, Kier et al. 2009, Negoita et al. 2016). All these mechanisms have deeply transformed insular and continental communities and therefore the degree to which they share species.

The theory of island biogeography (MacArthur and Wilson 1967) and empirical studies (Weigelt and Kreft 2013, Patiño et al. 2015) predict that immigration rates decrease with spatial distance. This implies that the number of species shared by islands and continents decrease with the distance between them. However, colonization is highly dependent on the abilities of species to disperse, which may be lineage specific. Consequently, some closely-related species may co-occur on spatially distant continental areas and islands, whereas other lineages may be spatially clustered and totally absent from remote places. For example, Patiño et al. (2015) demonstrated that seed plants species-richness decreased with island isolation whereas this was not the case for plants that produce spores, allowing dispersal on relatively longer distances. This dispersal filter makes that the effect of spatial distance is driven by the phylogenetic position of the species, hence blurring the expected effect of spatial distance on evolutionary history differences, *i.e.* phylogenetic divergence. Beyond spatial distance, environmental distance is another factor explaining why communities may share species or not (e.g. Nekola and White 1999, Tuomisto 2003, Carvajal-Endara et al. 2017). Different environmental conditions between ecosystems may prevent the settlement of some species because they cannot form viable populations in the environment to which they disperse. Again, the diversified responses of species to different environmental conditions may strongly reflect their position of species in the Tree of Life. Closely related species are expected to share traits and to occupy more similar niches thus co-occuring at places with similar environmental conditions (Webb et al. 2002). Following this reasoning, environmental distance would be a strong filter for the co-occurrence of closely related lineages and may increase phylogenetic divergence between communities (Tuomisto 2003, but see Gerhold et al. 2015). This may however not always be true. For instance, species from distinct lineages where trait evolution is predominantly convergent, as sometimes found in islands (Gillespie 2007), can co-occur in similar environments, even if they are over-dispersed in the phylogeny. On the contrary, very closely related species may adapt to highly diverging niches available to them when they arrive on an island (Evans et al. 2014). For all these reasons, the effect of spatial and environmental distance on phylogenetic divergence between islands and continents is still unclear (see Carvajal-Endara et al. 2017 for a regional example). A first objective of this study is thus to distinguish the contributions of spatial and environmental distance to phylogenetic divergence between insular and continental species pools.

However, spatial and environmental distances to the continent may only be part of the story explaining why species pools diverge between islands and continents and therefore why some islands are more divergent than others (Koenig et al. 2019). While we acknowledge that evolutionary and ecological dynamics in continents may have a role in explaining the phylogenetic divergence between islands and the mainland (Cronk 1992) here we focus on processes occurring in islands. The presence or absence of lineages on islands depend on ecological and evolutionary changes, resulting from colonization, diversification, extinctions that have a strong phylogenetic signature (Gillespie 2007). The importance of these processes in shaping island diversity may therefore be estimated by looking at the phylogenetic structure of communities (Weigelt et al. 2015, Carvajal-Endara et al. 2017). Nonetheless, analyses of community structure generally require the use of null models which, by randomizing species between pools, are largely dependent on the scale to which these pools are defined and make it difficult to detect the true community phylogenetic patterns (Graham and Fine 2008). Besides, as the insular pool mostly originate from mainland lineages, subjected to diversification or not, we assume here that additional processes could be revealed by looking at the phylogenetic divergence between insular and continental communities. Island-mainland comparison would be the “logical-way” to understand the evolution and assembly of the insular biota, but there are still very few studies which have taken this approach into consideration (Santos et al. 2016). Our second objective was thus to identify processes and estimate their contribution to the phylogenetic divergence between islands and continents. We assume that high divergence could be inferred from 1) long branches arising from relictualization or/and long-term survival of species in islands providing current and past refuge to plant species, 2) in situ speciation at the origin of evolutionary closely-related species in islands and of the share of deep tree branches between islands and continents, 3) low colonization events from the mainland so that only few branches would be shared, 4) high turnover leading to a high phylogenetic divergence both at the terminal and deep-branch level. To test these hypotheses we used measures of phylogenetic beta Mean Pairwise Distance (MPD_β_) - sensitive to variation at the deep branch level - and phylogenetic beta Mean Nearest Taxon Distance (MNTD_β_) - sensitive to variation at the terminal branch level - between insular and continental pools. Both of these indices of phylogenetic divergence were then linked to the phylogenetic structure of the species pool on islands and to island biophysical features, as both display traces of past mechanisms at the origin of island diversity.

We focused on Monocotyledons which is a very large clade, morphologically and functionally diverse, representing a quarter of flowering plant diversity. Monocots are distributed all across the globe and are well represented on islands. This, along with the existence of a well-resolved phylogeny and with database built by experts makes that Monocotyledon is a well-suited group to study the origins of phylogenetic divergence between insular and continental communities.

## Material and Methods

### Data

#### Islands and Continents

There is a wide diversity of islands types, but common properties are that they are isolated, well defined geographically and have well delimited boundaries. A total of 4,105 islands was used in this study. They were delimited with the Global Island Database provided by the United Nations Envioronment Programm (UNEP) (Depraetere and Dall 2007), and restricted to all isolated areas smaller than Australia occurring in oceans. Islands found within continental boundaries (e.g. in lakes, estuaries, rivers) were not considered. 610 continental areas were defined and delimited based one TDWG polygons at the 4th level (Brummitt et al. 2001).

#### Phylogeny

This study was based on a recent phylogeny of Monocots, which includes representatives of the great majority of genera (Tang et al. 2016). The use of four DNA regions (*rbcL, matK, ndhF* and nrITS DNA) allowed to place 1,816 genera on the phylogentic tree. The 710 genera for which genetic data were missing were included as polytomies within their respective clades using the most up-to-date taxonomy for each taxon (Tang et al. 2016).

#### Plant occurrences

Data on plant occurrences were obtained from the web platform of the Global Biodiversity Information Facility (GBIF). We downloaded all records, except fossil ones, and selected those intersecting the islands considered in this study. Then we used e-monocot database to *i.* control for synonyms and use only each species’ accepted name *ii.* to exclude non-native species occurrences *iii.* to control for species range by comparing occurrences from the GBIF data with those from e-monocot. E-monocot is a global database on monocotyledons which compiles occurrences assigned by experts to TDWG polygons. In addition, because aquatic plants tend to be highly evolutionary distant from all other species and might influence our estimates of phylogenetic divergence, we excluded all species belonging to the families Alismataceae, Acoraceae, Aponogetonaceae, Juncaceae, Juncaginaceae, Mayacaceae, Pontederiaceae, Potamogetonaceae, Ruppiaceae, Scheuchzeriaceae, Zosteraceae, Posidoniaceae, Cymodoceaceae and Hydrocharitaceae.

The dataset resulting from these filters comprised 2,568,386 occurrences representing 16,213 species and 1,562 genera in 4,105 islands, and 16,571,974 occurrences of 48,140 species from 2,178 genera in 610 continental areas.

### Estimates of phylogenetic divergence between islands and surrounding continents

#### Metrics

The phylogenetic divergence between insular and continental floras was calculated with phylogenetic Mean Pairwise Distance (MPD_β_) and Mean Nearest Taxon Distance (MNTD_β_), using the comdist function from R package picante (Kembel et al. 2010). MPD_β_ is the mean phylogenetic distance separating each pair of tips in two assemblages (here an insular and a continental pool). MPD_β_ estimates phylogenetic divergence in the global composition of islands and continents, and is sensitive to variation at the deep branch level (Webb et al. 2008) (Webb 2000). MNTD_β_ is the mean distance separating each tip in a site from its nearest relative occuring in another one and is sensitive to differences at the terminal branch level (Webb 2000, Webb et al. 2008). It is thus expected that these measures are complementary to the assessment of different processes conducting to the share of deep or short branches between islands and continents (e.g. an adaptative radiation following the settlement of a genera still present on the mainland will tend to decrease the number of shared short branches but will have a lesser influence on the number of shared deep branches).

#### Effect of spatial and environmental distance on phylogenetic divergence

We calculated MPD_β_ and MNTD_β_ between each island and each of the 10 nearest continental polygons. While we cannot be certain that the continental polygons defined here represent the true species pool for each island, it is likely that lineages present on islands originate from multiple relatively close continental regions. For example, Carvajal-Endara et al. (2017) estimated that the Galapagos species could have dispersed from 9 continental countries.. To test the sensitivity of the results to the number of continental polygons, we also measured MPD_β_ and MNTD_β_ between each island and each of the 5 and 20 nearest ones. We distinguished between oceanic and islands connected to the mainland during the Last Glacial Maximum – assumed to be a proxy for continental islands (Weigelt et al. 2013) – because different mechanisms may be at stake to explain divergence in each type of island (Whittaker and Fernández-Palacios 2007 but see Ali 2017).

As distance is assumed to be a main factor at differentiating biotas (53), we tested the predictive effect of spatial and environmental distances at explaining the variations in MPD_β_ and MNTD_β_ between insular and continental communities. Spatial distance was calculated in ArcGIS 10.3.1, as the distance between the centroids of an island and of a continental polygon. We then used four variables from (Fick and Hijmans 2017) reflecting distances in climatic conditions: differences in mean annual temperature, mean annual rainfall, mean annual solar radiation, mean annual wind speed. We did not include the differences of latitude and longitude due to colinearity with environmental variables, especially solar radiation and temperature, but their effect was tested separately (Appendix S1). All variables (e.g. difference in mean annual temperature) were then scaled and we estimated the average environmental distance as the euclidean distance among scaled variables (Tuomisto 2003). Finally, to account for other environmental dimensions not examined, we tested the effect of the number of different ecoregions between islands and continents (Olson et al. 2001).

We first used Generalized Linear Mixed Models (GLMM) to test the effect of spatial and environmental distances on MPD_β_ and MNTD_β_ between each island and a polygon in the continent. Second, we employed Boosted Regression Trees (BRT) (Elith et al. 2008) in order to make an informed decision about the relative contribution of each variable to phylogenetic divergence. BRT also displayed the direction of the relationship between the response and the predictors and we looked for interactions with the highest contribution (Elith et al. 2008). GLMM first tested the additive effects of all selected variables. We then used a second model in which all variables plus interaction between variables, identified thanks to the BRT method, were included. In GLMM, island’s identity was held as a random variable. Yet when BRT and GLMM methods led to contradictory results for a given variable, for example because of residual colinearity, we performed additional models, one for each technic, in which the variable in question was tested alone. Especially, this correction was needed to test the effect of euclidean environmental distance.

### Characterisation of the minimum divergence between islands and continents

To determine the divergence of each island with the most similar surrounding continental area and to further explore how mimimum divergence was distributed across the world we estimated its minimum MPD_β_ and MNTD_β_ values (minMPD_β_ and minMNTD_β_, respectively). We measured minMPD_β_ and minMNTD_β_ as the minimum values of divergence between an island and each of its 10 nearest continental polygons. Although insular species may have settled from multiple continental pools, we could not estimate the true original pool at the large scale we of our study, and we assumed that the minimum phylogenetic divergence from any continental polygon allowed to compare islands between them and to study the processes conducting to divergence between insular and continental pools.

#### Phylogenetic structure of insular communities

In order to explore how the phylogenetic structure of an insular community may influence minMPD_β_ and minMNTD_β_, we calculated 7 related attributes for each island: variance in pairwise diversity (VPD), mean pairwise distance (MPD_α_), the average and standard deviation in evolutionary distinctiveness (fair proportion index; ED), the proportion and the number of genera among the 10% of the most evolutionary distinct plants, and finally the number of genera in each island. Variance in pairwise diversity measures the regularity of the distribution of evolutionary history (Davies and Buckley 2012, Tucker et al. 2017). Evolutionary distinctiveness quantifies the number of relatives a species has and how phylogenetically distant they are (Faith 1992, Forest et al. 2007). MPD_α_ is the pairwise phylogenetic distance between the species of a given community (Webb 2000).

#### Abiotic correlates of phylogenetic divergence

Finally, we investigated which geographical, environmental, and historic factors, as well as sampling effort in islands may influence the minimum phylogenetic divergence with continents (Table 4). Note that these factors are intrinsic to an island, and that we did not use the difference in environmental condition between an island and the surrounding continents as in the previous section. We selected variables which had low colinearity with all others (Table 4), i.e. excluding variables with p>0.5 (Pearson correlation test). Latitude, likely to be an important factor of divergence representing a strong spatial effect, was tested independently due to its high colinearity with mean annual temperature. In addition, as our results may be biased by the fact that some islands were much less sampled than others, we re-ran the analysis without islands with low Incidence-based Coverage Estimate (ICE) values. ICE is calculated from the number of rare species in a sample and from species accumulation curves We calculated ICE by defining a sub-sample within an island as a set of observations obtained at a given date, and using the R function spp.est from the ‘Fossil’ package (Vavrek 2011). We then calculated the ICEr index as the ratio of the observed number of species in an area over the expected number of species estimated with ICE. The contribution and significance of distance, phylogenetic structure and abiotic factors was thus tested with a dataset by successively excluding islands with ICEr values lower than 0.2, 0.5 and 0.75. We found that results were not affected when these islands were discarded.

#### Modeling the influence of island community structure and abiotic factors on phylogenetic divergence

Using BRT and Generalized Linear Models (GLMs) we estimated the contribution and significance, respectively, of community structure and abiotic factors to minMPD_β_ and minMNTD_β_. No random variable was added in the models. Thanks to BRT we also estimated the strengths of interactions. The significance of the 2 interactions with the highest contributions were also tested with GLMs.

Exploring the relationship between abiotic features and community structure was not the main aim of this paper and was documented in previous studies (Weigelt et al. 2015, Carvajal-Endara et al. 2017), therefore the outcomes of this analyses is given as a supplementary material (Appendix S2).

## Results

### How spatial and environmental distance influence phylogenetic divergence between insular and continental communities

As estimated from boosted regression trees (BRT), variables with the highest importance at explaining phylogenetic beta mean pairwise distance (MPD_β_) were difference in solar radiation and difference in spatial distance (Table 2, Appendix S3). Difference in rainfall also had a high contribution in the case of oceanic islands. When accounting for island identity as a random effect in generalized linear mixed models (GLMMs), MPD_β_ significantly increased with spatial distance and rainfall, regardless if islands were continental or oceanic. Differences in solar radiation and temperature had an unexpected negative relationship with MPD_β_, probably reflecting their non-linear relationships with the difference in latitude in the tropics (Appendix S1). Indeed, when tropical islands were excluded, the effect of these two variables had a significant positive relationship with MPD_β_. The remaining variables (wind speed, number of ecoregions and Euclidean environmental distance) had lower contributions but significant positive relationship with MPD_β_ (Table 2; Appendix S3). Looking at MNTD_β_, variables which generally had a high contribution were the difference in annual solar radiation, spatial distance and the euclidean environmental distance. However, contribution of spatial distance was much lower in continental islands (Table 2, Appendix S3). When island identity was integrated as a random effect in GLMMs all variables had a positive relationship with MNTD_β_. Overall, the choice of number of continental polygons used to measure divergence did not change the major results.

**Table 1.** The 15 abiotic variables tested and sources of data

| *Variable* | *Unit* | *Source* |
| --- | --- | --- |
| ***Geographical variables*** |  |  |
| *Latitude* | Decimal degrees | Depraetere & Dall (2007) |
| *Longitude* | Decimal degrees | Depraetere & Dall (2007) |
| *Minimum distance to continent* | km | Depraetere & Dall (2007) |
| *Proportion of surrounding land mass (SLMP)* |  | Weigelt et al. (2013) |
| *Area* | km<sup>2</sup> | Depraetere & Dall (2007) |
| *Elevation* | m | Weigelt et al. (2013),<br>Depraetere & Dall (2007) |
| ***Climate*** |  |  |
| *Number of ecosystems* |  | Olson et al. (2001) |
| *Mean annual temperature* | °C | Fick & Hijmans, (2017) |
| *Mean annual rainfall* | mm | Fick & Hijmans, (2017) |
| *Rainfall seasonality* | mm | Fick & Hijmans, (2017) |
| *Mean annual wind speed* | m.s<sup>-1</sup> | Fick & Hijmans, (2017) |
| *Standard deviation of vapor pressure* | kPa | Fick & Hijmans, (2017) |
| ***Historical factors*** |  |  |
| *Connection to the mainland during the LGM (GMMC)* |  | Weigelt et al. (2013) |
| *Velocity of past climate change* | y.m<sup>-1</sup> | Sandel et al. (2011) |
| ***Sampling effort*** |  |  |
| *ICE<sub>r</sub>* |  | Lee & Chao (1994) |

**Table 2.**
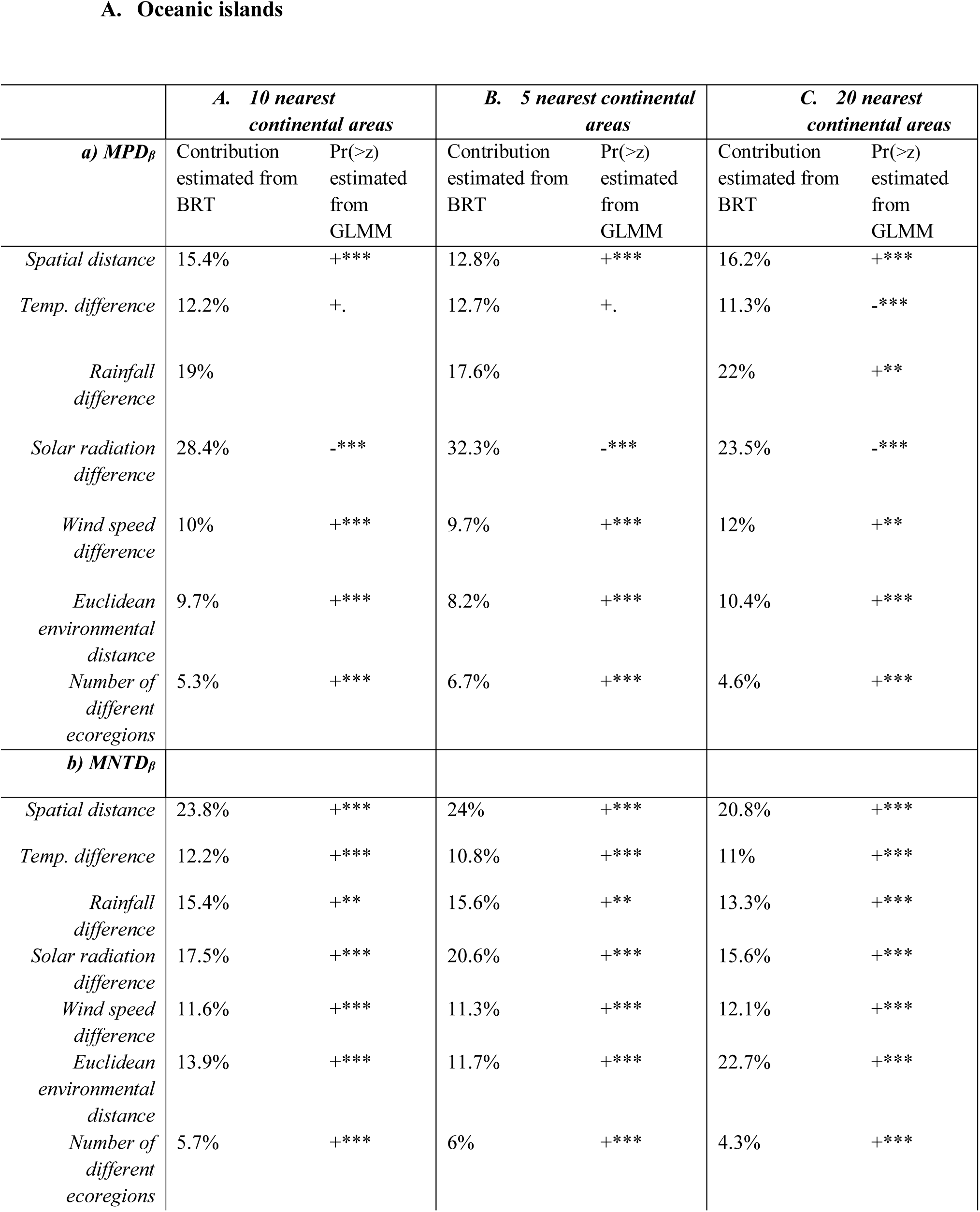
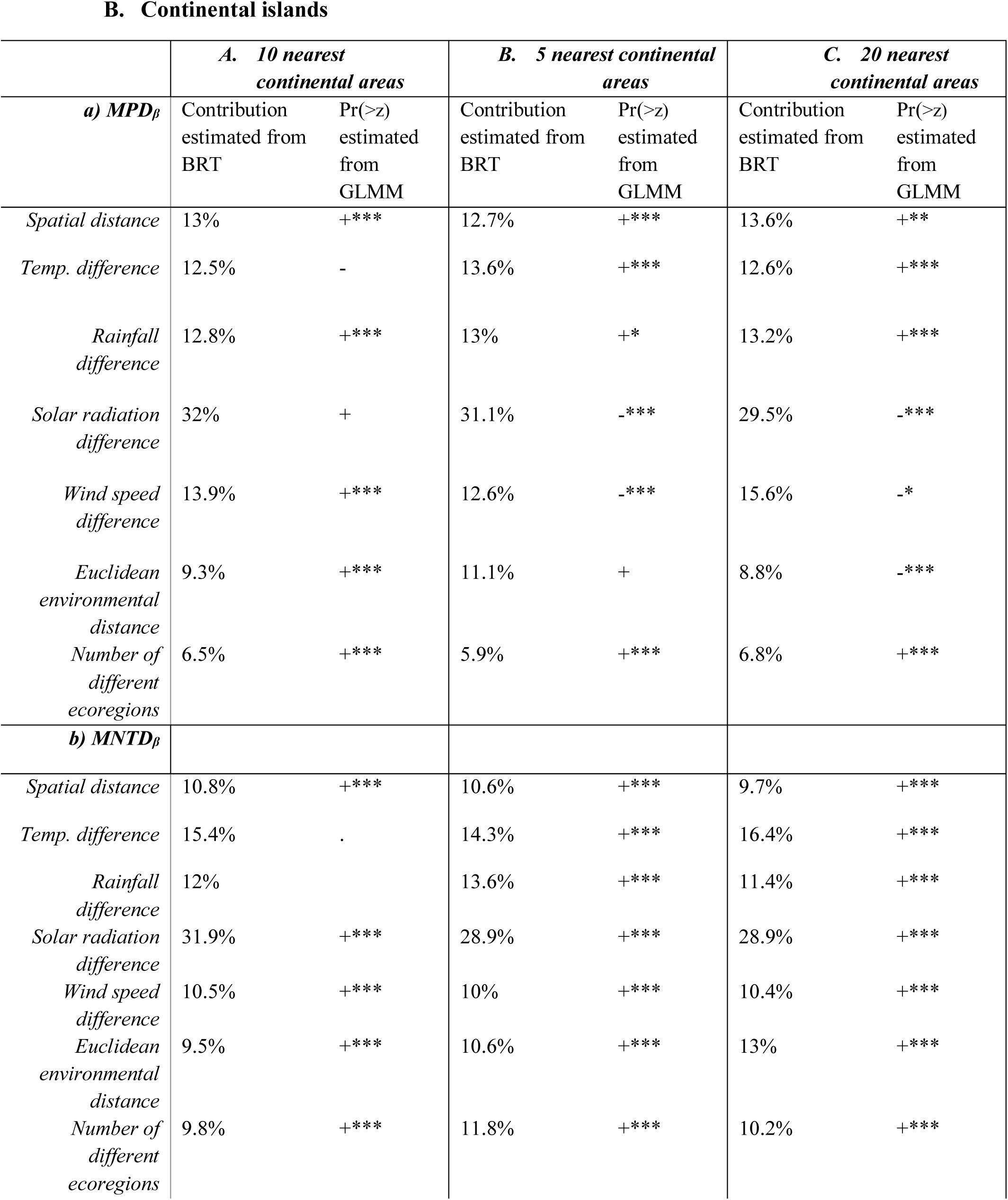
Effects of spatial and environmental distance on MPD**β** and MNTD**β** between islands and continents measured from BRT and GLM: “***” indicate Pr(>z) <0.0001; “**” indicate Pr(>z) <0.001; “*” indicate Pr(>z) <0.01 “.” indicate Pr(>z) <0.1; + and – indicate the direction of the relationship that is positive or negative, respectively. Column A presents the observed results. Column B and C represents data used as a sensitivity analysis.

### Spatial patterns of island-continent divergence

Islands with floras highly divergent from continents when using minMPD_β_ were found worldwide, even though minMPD_β_ values tended to be the highest around the equator (figure 2a). Madagascar, Fiji, Indonesia and West Indies are exemples of islands with high minMPD_β_ (figure 2a, Appendix S4).

Spatial patterns of minMNTD_β_ roughly differed from those of minMPD_β_. Especially, there were relatively more islands with high minMNTD_β_ in the Southern hemisphere. Yet, when looking at extreme values, islands with the highest minMNTD_β_ were similar to those having the highest values minMPD_β_. Exemples are Madagascar, Fiji, Indonesia, Cook Islands and American Samoa (figure 2b, Appendix S4).

**Figure 1.**
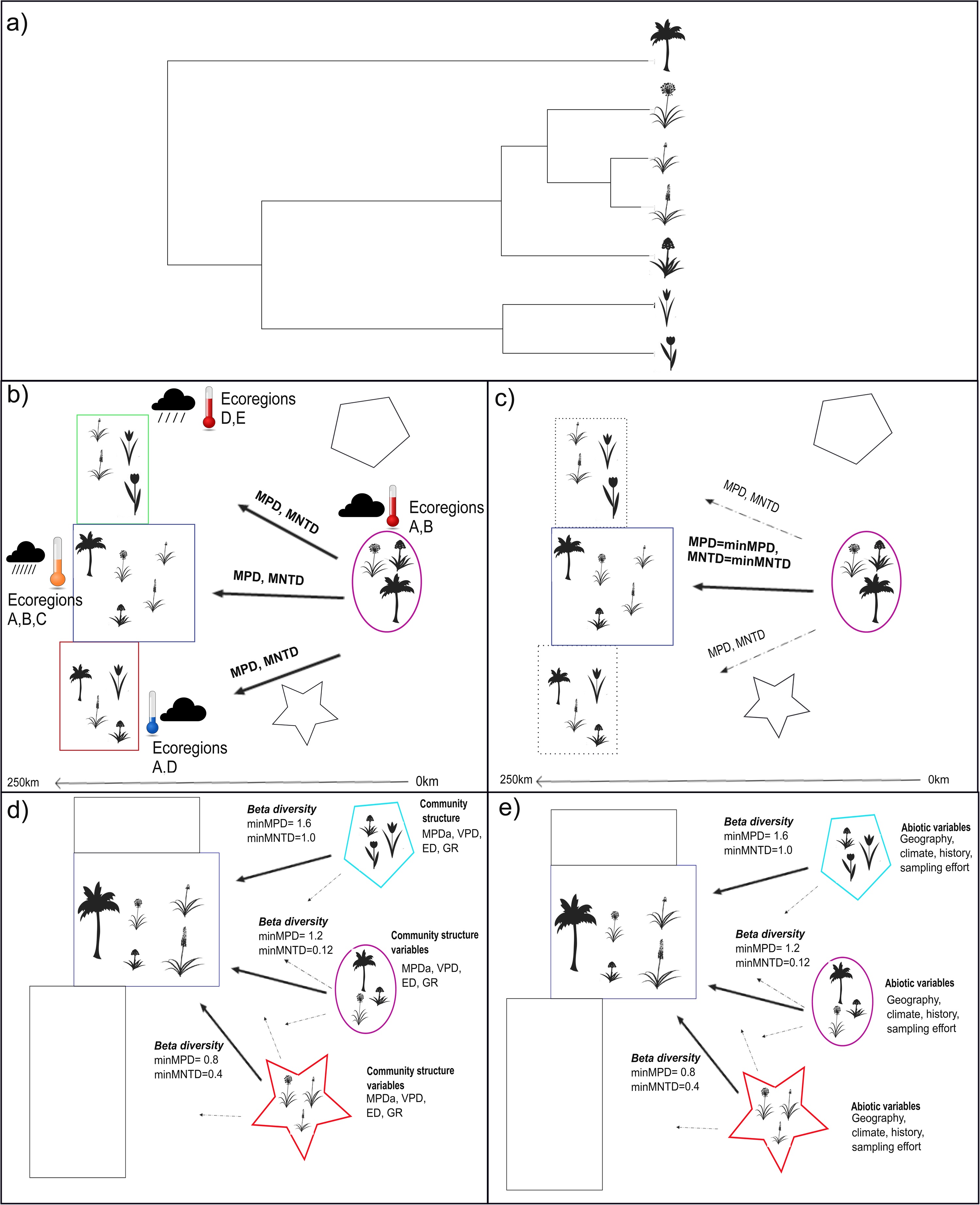
Methodological procedure a) Figure of a phylogenetic tree b) Beta diversity between an islands and the surrounding continents. MPD and MNTD were estimated from the 5, 10, 20 closest continental areas to each island as well as from all continents belonging to a similar biome as each island. Relationships between MPD and MNTD with spatial and environmental distances c) Identification of minimum beta diversity values for each island and each index (MPDmin and MNTDmin). d) Relationships minimum between beta diversity and community structure measures (see also table 2). *MPDa= alpha Mean pairwise Distance; VPD= Variance in Pairwise Distance* *ED= Evolutionary Distinctiveness; GR= Genus Richness* e) Relationships between minimum beta diversity and abiotic factors (see also table 3)

**Figure 2.**
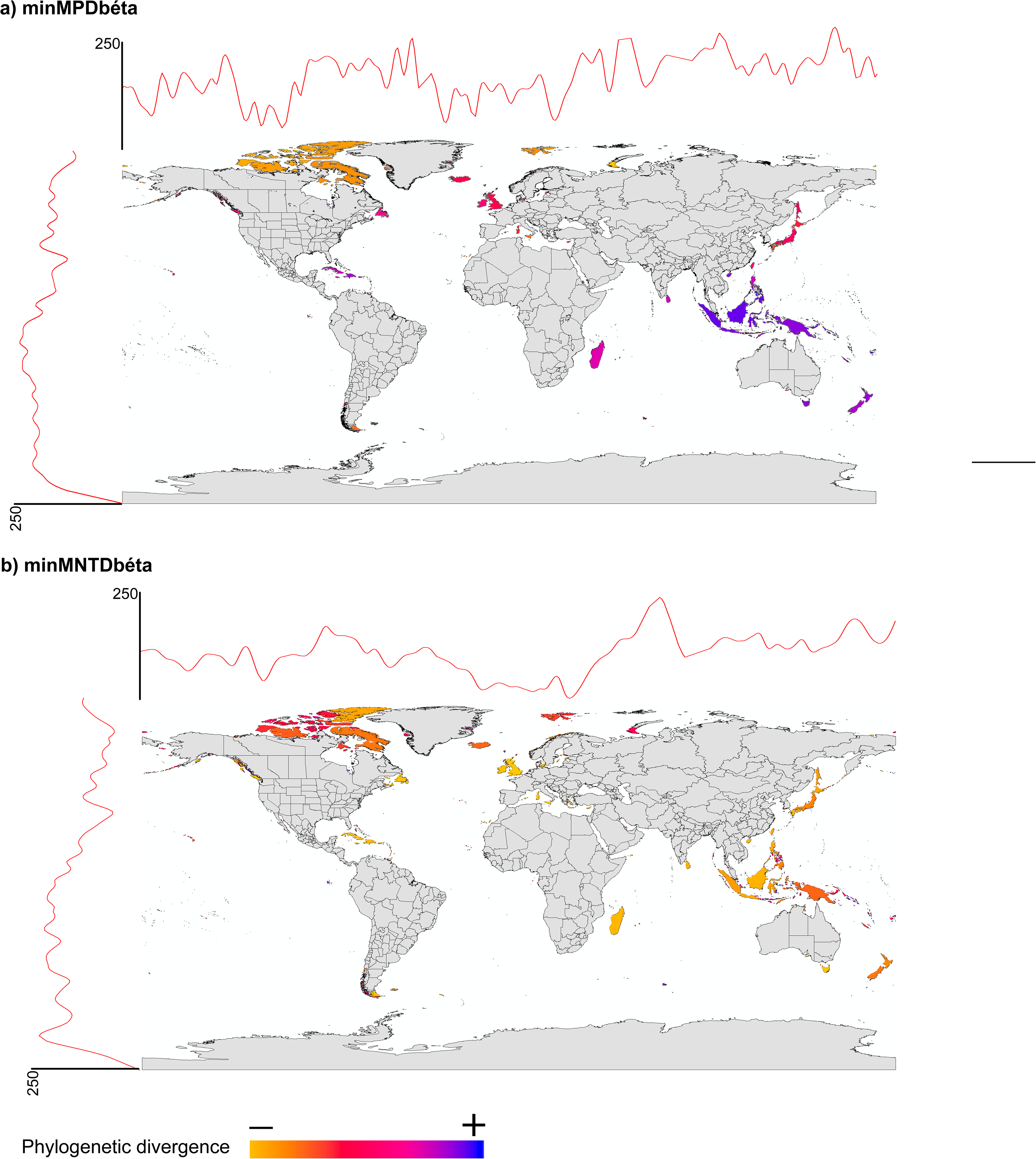
Spatial distribution of the minimum phylogenetic divergence between islands and the 10 nearest continental areas for a) Mean Pairwise Distance b) Mean Nearest Taxon Distance.

### Effect of phylogenetic structure on phylogenetic divergence between island and continents

We found that the effect of insular community structure varied depending on whether minMNTD_β_ or minMPD_β_ was used. The average Evolutionary Distinctiveness (ED) of insular monocots had the highest contribution and a significant positive effect on minMPD_β_ (table 3). On the contrary, standard deviation in ED had a significant negative effect. MPD_α_ and genus richness ranked second and fifth regarding their contribution to minMPD_β_ (contributions equal to 31.8% and 2.1%, repectively). Both variables displayed a significant positive relationship with minMPD_β_. The highest contribution to minMNTD_β_ was the number of insular genera (contribution = 42.3%), which had a significant negative effect. Other variables with a relatively high contribution were VPD (contibution=30.6%) and MPD_α_ (contribution=15.04%), both having a significant negative relationship with minMNTD_β_.

**Table 3.**
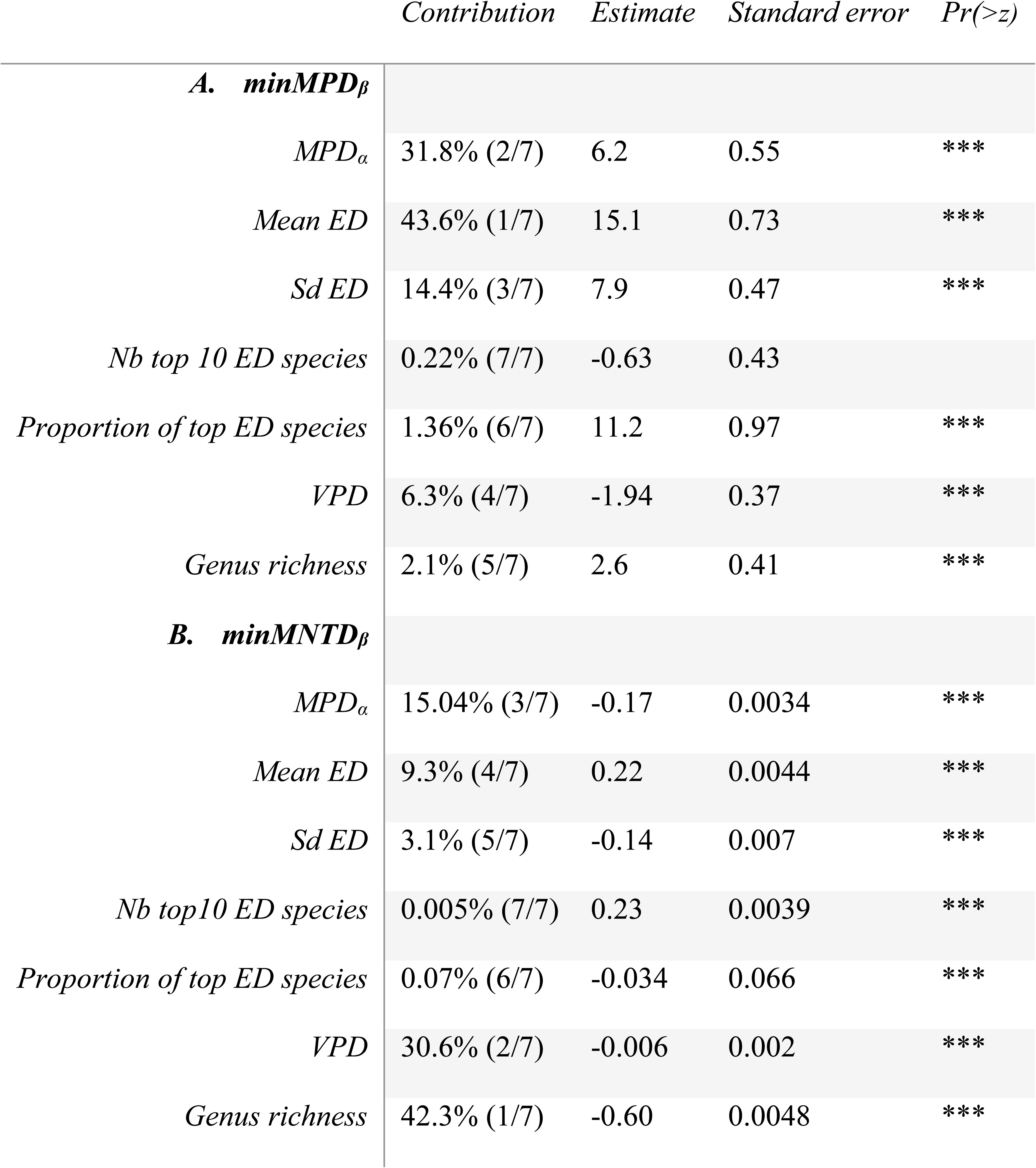
Contribution, direction and significance of variables of phylogenetic structure to a) minimum MPD _β_ and b) minimum MNTD _β_ between islands and continents.

**Table 4:** Contribution, direction and significance of abiotic variables to a) minimum MPD _β_ between islands and continents and b) minimum MNTD _β_ betweeen islands and continents.

A. $\min MPD_{\beta}$
|  |  | *Contribution to*<br>*$\min MPD_{\beta}$* | *Estimate* | *Standard*<br>*error* | *Pr(>z)* |
| --- | --- | --- | --- | --- | --- |
| *Geographic*<br>*variables* | Area | 1.6% (11/15) | -1.2 | 0.58 | * |
|  | SLMP | 1.5% (12/15) | 1.1 | 0.86 |  |
|  | Elevation | 0.9% (13/15) | -0.17 | 0.68 |  |
|  | Minimum distance to<br>continents | 5.8% (8/15) | 1.2 | 0.54 | * |
|  | Latitude | 35.5% (1/15) | -7.9 | 0.87 | *** |
|  | Longitude | 9.8% (3/15) | 0.17 | 0.55 |  |
| *Climatic*<br>*variables* | Number of ecosystems | 0.2% (15/15) | 1.8 | 0.56 | ** |
|  | Mean annual temperature | 9.1% (4/15) | 7.3 | 0.94 | *** |
|  | Mean annual rainfall | 10.7% (2/15) | 5.06 | 0.68 | *** |
|  | Rainfall seasonality | 7.7% (5/15) | -6.3 | 0.65 | *** |
|  | Mean annual wind speed | 6.3% (6/15) | -5.0 | 0.75 | *** |
|  | Standard deviation in vapour<br>pressure | 5.9% (7/15) | -3.6 | 0.63 | *** |
| *Historical*<br>*variables* | Velocity of past climate<br>change | 4.5% (9/15) | 4.9 | 0.72 | *** |
|  | GMMC | 0.2% (14/15) | 7.7 | 1.62 | *** |
| *Sampling effort* | ICEr | 1.7% (10/15) | -0.31 | 0.48 |  |
| *Interactions* | Latitude:Longitude |  | -6.5 | 0.53 | *** |
|  | Longitude:Temperature |  | 2.6 | 0.54 | *** |

B. $\min MNTD_{\beta}$
|  |  | *Contribution to*<br>*$\min MNTD_{\beta}$* | *Estimate* | *Standard*<br>*error* | *Pr(>z)* |
| --- | --- | --- | --- | --- | --- |
| *Geographic*<br>*variables* | Area | 4.3% (6/15) | 0.034 | 0.003 | *** |
|  | SLMP | 2.6% (9/15) | -0.098 | 0.004 | *** |
|  | Elevation | 1.6% (13/15) | -0.12 | 0.0039 | *** |
|  | Minimum distance to continent | 1.8% (10/15) | 0.045 | 0.0032 | *** |
|  | Latitude | 31.1% (1/15) | -0.22 | 0.004 | *** |
|  | Longitude | 9.7% (3/15) | -0.07 | 0.0029 | *** |
| *Climatic variables* | Number of ecosystems | 0.8% (14/15) | -0.063 | 0.0037 | *** |
|  | Mean annual temperature | 4.3% (5/15) | 0.017 | 0.0049 | *** |
|  | Mean annual rainfall | 3.1% (7/15) | 0.066 | 0.0033 | *** |
|  | Rainfall seasonality | 1.7% (11/15) | 0.037 | 0.0035 | *** |
|  | Mean annual wind speed | 1.6% (12/15) | -0.08 | 0.0039 | *** |
|  | Standard deviation in vapour pressure | 7.0% (4/15) | -0.004 | 0.006 | . |
| *Historical variables* | Velocity of past climate change | 2.7% (8/15) | 0.07 | 0.0036 | *** |
|  | GMMC | 0.03% (15/15) | -0.058 | 0.009 | *** |
| *Sampling effort* | ICEr | 26.7% (2/15) | -0.10 | 0.0031 | *** |
| *Interactions* | Latitude:Longitude |  | -0.14 | 0.0023 | *** |
|  | ICE:Area |  | -0.036 | 0.0027 | *** |

### Abiotic correlates of phylogenetic divergence

Latitude was the variable with highest contribution to explain minMPD_β_ and its effect was significantly negative. Interaction of latitude and longitude also had a significant negative effect: effect of low latitude on phylogenetic divergence between islands and continents was stonger at high longitude. Other geographic variables had a low contribution to minMPD_β_ such as the distance to the nearest continental polygon and area which had slightly significant positive and negative relationships, respectively. Climatic variables generally had a moderate contribution to minMPD_β_. Specifically mean annual rainfall and temperature had the highest contributions among all abiotic variables (Table 4a). Climatic variables all had a significant effect on minMPD_β_. It was positive for mean annual temperature, mean annual rainfall and the number of ecosystems, but negative regarding rainfall seasonality, annual wind speed, and standard deviation in vapour pressure. Although historical factors had a relatively low contribution to MPD_β_, we found a significant positive effect of the velocity of past climate change and of the past connexion to the mainland.

Regarding minMNTD_β_ between islands and continents, latitude (contribution=31.1%), the Relative Incidence-based Coverage Estimator (ICEr; contribution=26.7%) and longitude (contribution=9.7%) were important to explain minMNTD_β_, with a significant negative effect (Table 4b). ICE_r_ displayed a significant negative interaction with area, meaning that poorly sampled and small islands had the highest minMNTD_β_ values. Contribution of environmental variables was low. Among them, mean annual temperature (contribution=4.3%; significant positive effect) and standard deviation in vapour pressure (contribution=7.0%; significant negative effect) had the highest importance. As for historical variables, velocity of past climate change had a significant positive relationship on minMNTD_β_, whereas past connection to the mainland, contrary to what was observed for minMPD_β_, had a significant negative effect (Table 4b, Fig S2).

## Discussion

Difference in species composition between islands and continents has already been well-documented. This study goes one step further by showing that insular and continental pools also differ in their evolutionary history and that the degree to which they share phylogenetic branches depend on the phylogentic signature of the processes shaping diversity.

### a. Divergence due to spatial and environmental distance

While effect of the distance to the continent is recognised to be a main driver of species richness and community composition in islands (MacArthur and Wilson 1967, Brown and Kodric-Brown 1977, Rosindell and Phillimore 2011, Cabral et al. 2014), the present study gives evidence that it is also true at the phylogenetic level. This result was expected because a difference in species composition also means that the phylogenetic branches supporting these species are distinct between pools. As spatial distance acts as a dispersal filter reducing the number of individuals exchanged between continents and islands, it also reduces the number of shared phylogenetic branches between them. More specifically, our results suggest that among the insular species and phylogenetic branches originating from the mainland, most of them are or were present on the nearest continents. One of the main benefits of using phylogenetic approaches is that it allows investigating the mechanisms of dispersal filtering that have a strong phylogenetic signature. For instance, some studies show that dispersal toward some islands is lineage specific due to phylogenetically clustered traits such as seed size. However, we found that distance had a similar influence on the co-occurrence of deep or terminal branches indicating that a dispersal filter towards species clustered in the phylogeny is not detected at a global scale.

Distance to the continent is only the first filter of species settlement, although some individuals may reach an island, only a few species may truly establish themselves due to specific niche requirements. Several studies showed the prevailance of environmental over dispersal filtering (Tuomisto 2003, Carvajal-Endara et al. 2017). However, at a global scale it was unclear which of those filters had the strongest influence on the divergence between islands and continents. Strong environmental filtering is expected to generate species phylogenetic clustering due to phylogenetic trait conservatism (Webb et al. 2002). Despite the fact that distinct environmental conditions reduce the number of shared branches between islands and continents, the genera they supported were likely not clustered the phylogeny (Table 1).

In the case of continental islands, increased divergence with environmental distance may result from two complementary factors. On the one hand, lineages that were present at the island birth and that persist nowadays may have survived thanks to the continuity of a stable island climate through time (Sandel et al. 2011, Weigelt and Kreft 2013). On the opposite, we showed that high velocity of past climate change result in high phylogenetic divergence between continents and islands.

This, with the complex geological history following the breakup of continental islands, the presence of a complete set of species at birth and more frequent arrivals may also explain the lesser importance of spatial distance on divergence when compared to oceanic islands.

### b. How island features and community structure drive phylogenetic divergence

Distance is a key determinant of species arrival and establishment on islands but further ecological and evolutionary processes are at the origin of divergence between islands and continents. The study of phylogenetic divergence may allow distinguishing several of them. Geological and climatic history of islands drive the long-term survival of species promoting their evolutionary distinctiveness (ED). The monocotyledons with highest ED values, usually hold long branches and have few close relatives. So, ED results either from ancient speciation events that gave rise to a single of few species, or from the evolutionary isolation of a species through the extinction of its nearest-relatives, i.e. relictualization (Gillespie 2007, Grandcolas and Trewick 2016). We found that the more evolutionary distinct the monocotyledon on an island the more divergent from continental pools they were, but that ED had a lower contribution on the divergence of the most closely-related taxa rather than on the divergence beween all taxa. This is likely explained because many lineages with high ED may still be present on the continent (Jetz et al. 2014). Indeed, when high ED monocotyledons are present on both continents and islands, the phylogenetic distance may remain high with all other taxa, but may be lower when it is estimated only from the most closely-related taxa. The present results thus indicate that relictualiz:ation is likely to have been less frequent than evolutionary processes in explaining the origin of ED monocotyledons. Besides, we also observed that islands where mean ED is high were generally connected to the continent in the past. Insular monocotyledons which are currently evolutionary distinct may have therefore probably appeared on the mainland before islands became fragmented.

Phylogenetic divergence is not only due to ancient and isolated lineages, but also to recent and clustered taxa absent on the mainland. Our results show that when phylogenetic distance of the insular pool is low and regular, island and mainland flora highly diverge. At the generic level, this may correspond to islands where environmental, dispersal, or biotic filters have allowed a majority of closely related monocot lineages to establish (Cadotte and Tucker 2017, Koenig et al. 2019). However, as we did not find an effect of environmental distance or spatial distance on the occurrence of closely-related genera on islands, additional factors potentially causing such clustered phylogenetic structure should be investigated. At the species level it may be expected that regular and low phylogenetic distance may cause high phylogenetic divergence due to evolutionary radiations. Recent species radiations may form clusters of endemic species with regular and low phylogenetic distance, conducting to few terminal branches shared with the mainland. Such radiations are more likely to occur on remote oceanic islands. Indeed, in these islands, immigration followed by in-situ speciation is the main process at the origin of species diversity: colonization rate is so slow that the main processes to fill the niche space are evolution or adaptation (Gillespie 2007, Emerson and Gillespie 2008).

Besides diversification and extinction, colonization is likely to be another important cause of phylogenetic divergence. Colonization from the continent may increase the co-occurence of similar lineages between islands and continents. We found that three factors related to high colonization rates increase phylogenetic similarity between island and continental pools, confirming our results about the effect of dispersal filtering: small spatial distance from the continent, large propotion of surrounding land mass and high wind speed. Proximity to the continent, and more generally to any other land mass, facilitates exchanges and thus high phylogenetic similarity at both the terminal and the deep branch levels (MacArthur and Wilson 1967). It probably means that genera having established on these islands are neither over-dispersed nor clustered in the phylogeny. Regarding wind speed, its negative relationship with divergence highlights that monocot dispersal capacities may have a critical role to explain colonization success. The importance of wind speed to divergence tends to be stronger at deep than at terminal branch level. Genera whose representatives can disperse through anemochory may consequently tend to be over-dispersed in the phylogeny but functional trait analyses are needed to test this assumption. Plants may also have reached islands thanks to animals, ocean currents, or human conveyance. To understand how this mode of dispersal influences which lineages co-occur on islands and continents, future research may look at the dispersal traits of lineages having colonized islands and at the directionality and trajectories of dispersal factors (Patiño et al. 2015, Gillespie et al. 2012).

Other key factors of divergence between islands and continents are species turnover and speciation rates. Islands with high temperature and intense but relatively regular precipitations, i.e. islands with a tropical rainforest climate, were more likely to harbor a flora that diverges from the mainland flora. Rainfall and temperature, but also vapour pressure, are related to water and energy supply, which may increase turnover but also speciation rates, conducting to divergent communities from the mainland (Hawkins et al. 2003, Beres et al. 2008). While speciation are more frequent at the species level, it also occurred in monocots at the generic level, especially in the tropics. This divergence was important to both deep and short branches, which suggests that turnover is not clustered in the phylogeny. High energy and water supply is common in the tropics and around the equator, but these are also places where many processes departing from current environment, such as evolutionary, climatic and geologic history, have let a strong imprint on species diversity and divergence. Mechanisms at the origin of this latitudinal gradient in species diversity still fuels the debate and consequences in terms of divergence between species pools remain to be investigated.

Finally, not only current but also past environmental conditions are important to explain phylogenetic divergence, in particular on continental islands. Velocity of past climate change may actually be associated to low environmental stability (Fjeldsä 1994) and past species extinctions of insular species (Sandel et al. 2011) but also to strong environmental filtering conducting to dissimilar flora between continents and islands

### Directions for future research

Although we reduced biases in occurrence data through different methods, working at a global scale implied that the dataset can be subject to sampling biases over all islands, due to varying priorities along data acquisition history. Especially, more intense sampling in some less studied islands, but also the mobilisation of yet non-digitized herbarium data will certainly lead to higher robustness of the results obtained here. In addition, although the use of the generic level and of presence only data have been important in reducing biases in monocot spatial distribution, this may have prevented from detecting the importance of some mechanisms on divergence, for example adaptative radiations.

A second point that may stimulate further research is that we have only been interested in abiotic factors to explain divergence between insular and continental flora. Biotic factors, especially competition and interactions conducting to co-evolution, may act as a biotic filter, shape the phylogenetic structure of communities and their divergence with the mainland (Wilson 1969, Webb et al. 2002, Gillespie 2007, Weigelt et al. 2015). We therefore encourage to direct future research on the role of biotic factors in shaping the phylogenetic structure of communities and how they may explain divergence between insular and continental pools.

Moreover, at considering the distance from each island to the nearest continent we ignored potential dispersion between islands and especially the role of island-hopping which may be an important process in the composition of insular communities (Sillero et al. 2018). Insular species diversity has also a strong temporal dimension: age reflects time for speciation and colonization (Emerson and Gillespie 2008), area but also past distance to the continent varied through geological times (Norder et al. 2018) and are expected to have a strong influence on patterns of beta diversity. In addition the biological classification in a simple dichotomy, i.e. oceanic versus continental islands, may be complemented by introducing other island types (Ali 2017).

### Conclusion: new insights on plant community divergence between island and continents thanks to phylogenetic approaches

Phylogenetic approaches were rarely employed to understand patterns and origin of plant diversity in islands (Kreft and Jetz 2007, Kreft et al. 2008, Kier et al. 2009, but see Weigelt et al. 2015). Here, using approaches based on phylogenetic divergence, we provided new insights on the continental origin of plant diversity in islands. At a global scale spatial and environmental distance act as filters to species establishment on islands although they are not directed toward specific lineages. On continental islands, an unchanged climate may have allowed species to persist following the isolation process. We then investigated the importance of island features and phylogenetic structure in explaining deep or short branches shared by islands and the mainland. This showed that reasons for phylogenetic divergence were *i.* the persistence of evolutionary original species *ii.* regular and low phylogentic distance among species caused by - for example - speciation in remote archipelagos *iii.* colonization of non-clustered taxa *iv.* high turnover and speciation rates. We further encourage future use of phylogenetic approaches in island biogeography, for example to disentangle the role of age or of biotic factors on patterns of island diversity.

## Supporting information

Supplementary result 1

Supplementary result 2

Supplementary result 3

## Appendix legends

**Appendix S1**. Analysis of the relationship between solar radiation, latitude and phylogenetic divergence

**Appendix S2**. Effect of island features on phylogenetic structure of insular pools

**Appendix S3**. Boosted Regression Tree contributions of spatial and environmental distance to phylogenetic divergence

## Data

Data will be made available on the Dryad repository

## Declarations

### Funding

This study has been supported by the French State through the Research National Agency under the LabEx ANR-10-LABX-0003-BCDiv, within the framework of the program ‘Investing for the future’ (ANR-11-IDEX-0004-02).

### Author contributions

The first author is the major contributor to this study. The last co-author is the leading co-author of this work

## References

Ali, J. R. 2017. Islands as biological substrates: classification of the biological assemblage components and the physical island types. - J. Biogeogr. 44: 984–994.

Beres, K. A. et al. 2008. Rotifers and Hubbell’s unified neutral theory of biodiversity and biogeography. - Nat. Resour. Model. 18: 363–376.

Brown, J. H. and Kodric-Brown, A. 1977. Turnover Rates in Insular Biogeography: Effect of Immigration on Extinction. - Ecology 58: 445–449.

Brummitt, R. et al. 2001. World geographical scheme for recording plant distributions.

Cabral, J. S. et al. 2014. Biogeographic, climatic and spatial drivers differentially affect α, β and γ diversities on oceanic archipelagos. - Proc. R. Soc. B Biol. Sci. 281: 20133246–20133246.

Cadotte, M. W. and Tucker, C. M. 2017. Should Environmental Filtering be Abandoned? - Trends Ecol. Evol. 32: 429–437.

Carvajal-Endara, S. et al. 2017. Habitat filtering not dispersal limitation shapes oceanic island floras: species assembly of the Galápagos archipelago. - Ecol. Lett. 20: 495–504.

Cronk, Q. C. B. 1992. Relict floras of Atlantic islands: patterns assessed. - Biol. J. Linn. Soc. 46: 91–103.

Davies, T. J. and Buckley, L. B. 2012. Exploring the phylogenetic history of mammal species richness: A phylogenetic history of mammal richness. - Glob. Ecol. Biogeogr. 21: 1096–1105.

Depraetere, C. and Dall, A. 2007. IBPoW Database. A technical note on a global dataset of islands.: 58.

Elith, J. et al. 2008. A working guide to boosted regression trees. - J. Anim. Ecol. 77: 802–813.

Emerson, B. C. and Gillespie, R. G. 2008. Phylogenetic analysis of community assembly and structure over space and time. - Trends Ecol. Evol. 23: 619–630.

Evans, M. et al. 2014. Insights on the Evolution of Plant Succulence from a Remarkable Radiation in Madagascar (Euphorbia). - Syst. Biol. 63: 697–711.

Faith, D. P. 1992. Conservation evaluation and phylogenetic diversity. - Biol. Conserv. 61: 1–10.

Fick, S. E. and Hijmans, R. J. 2017. WorldClim 2: new 1-km spatial resolution climate surfaces for global land areas: new climate surfaces for global land areas. - Int. J. Climatol. 37: 4302–4315.

Forest, F. et al. 2007. Preserving the evolutionary potential of floras in biodiversity hotspots. - Nature 445: 757–760.

Gerhold, P. et al. 2015. Phylogenetic patterns are not proxies of community assembly mechanisms (they are far better). - Funct. Ecol. 29: 600–614.

Gillespie, R. G. 2007. Oceanic islands: models of diversity. - Encycl. Biodivers.: 1–13.

Gillespie, R. G. et al. 2012. Long-distance dispersal: a framework for hypothesis testing. - Trends Ecol. Evol. 27: 47–56.

Graham, C. H. and Fine, P. V. A. 2008. Phylogenetic beta diversity: linking ecological and evolutionary processes across space in time. - Ecol. Lett. 11: 1265–1277.

Grandcolas, P. and Trewick, S. A. 2016. What Is the Meaning of Extreme Phylogenetic Diversity? The Case of Phylogenetic Relict Species. - In: Pellens, R. and Grandcolas, P. (eds), Biodiversity Conservation and Phylogenetic Systematics: Preserving our evolutionary heritage in an extinction crisis. Springer International Publishing, pp. 99–115.

Hawkins, B. A. et al. 2003. Energy, water, and broad-scale geographic patterns of species richness. - Ecology 84: 3105–3117.

Jetz, W. et al. 2014. Global Distribution and Conservation of Evolutionary Distinctness in Birds. - Curr. Biol. 24: 919–930.

Kembel, S. W. et al. 2010. Picante: R tools for integrating phylogenies and ecology. - Bioinformatics 26: 1463–1464.

Kier, G. et al. 2009. A global assessment of endemism and species richness across island and mainland regions. - Proc. Natl. Acad. Sci. 106: 9322–9327.

Koenig, C. et al. 2019. Disharmony of the world’s island floras: Supporting Information. - bioRxiv in press.

Kreft, H. and Jetz, W. 2007. Global patterns and determinants of vascular plant diversity. - Proc. Natl. Acad. Sci. 104: 5925–5930.

Kreft, H. et al. 2008. Global diversity of island floras from a macroecological perspective. - Ecol. Lett. 11: 116–127.

MacArthur, R. H. and Wilson, E. O. 1967. The theory of island biogeography. - Princeton, NJ: Princeton University Press.

Negoita, L. et al. 2016. Isolation-driven functional assembly of plant communities on islands. - Ecography 39: 1066–1077.

Nekola, J. C. and White, P. S. 1999. The distance decay of similarity in biogeography and ecology. - J. Biogeogr. 26: 867–878.

Norder, S. J. et al. 2018. A global spatially explicit database of changes in island palaeo-area and archipelago configuration during the late Quaternary. - Glob. Ecol. Biogeogr. 27: 500–505.

Olson, D. M. et al. 2001. Terrestrial Ecoregions of the World: A New Map of Life on Earth. - BioScience 51: 933.

Patiño, J. et al. 2015. Island floras are not necessarily more species poor than continental ones. - J. Biogeogr. 42: 8–10.

Patiño, J. et al. 2017. A roadmap for island biology: 50 fundamental questions after 50 years of *The Theory of Island Biogeography*. - J. Biogeogr. 44: 963–983.

Rosindell, J. and Phillimore, A. B. 2011. A unified model of island biogeography sheds light on the zone of radiation: A unified model of island biogeography. - Ecol. Lett. 14: 552–560.

Sandel, B. et al. 2011. The Influence of Late Quaternary Climate-Change Velocity on Species Endemism. - Science 334: 660–664.

Santos, A. M. C. et al. 2016. New directions in island biogeography: New directions in island biogeography. - Glob. Ecol. Biogeogr. 25: 751–768.

Sillero, N. et al. 2018. Analysing the importance of stepping-stone islands in maintaining structural connectivity and endemicity. - Biol. J. Linn. Soc. 124: 113–125.

Steinbauer, M. J. et al. 2016. Biogeographic ranges do not support niche theory in radiating Canary Island plant clades: Niche theory in radiating Canary Island plant clades. - Glob. Ecol. Biogeogr. 25: 792–804.

Tang, C. Q. et al. 2016. Global monocot diversification: geography explains variation in species richness better than environment or biology. - Bot. J. Linn. Soc. in press.

Tucker, C. M. et al. 2017. A guide to phylogenetic metrics for conservation, community ecology and macroecology: A guide to phylogenetic metrics for ecology. - Biol. Rev. 92: 698–715.

Tuomisto, H. 2003. Dispersal, Environment, and Floristic Variation of Western Amazonian Forests. - Science 299: 241–244.

Vavrek, M. J. 2011. Fossil: palaeoecological and palaeogeographical analysis tools. - Palaeontol. Electron.

Wallace, A. R. 1880. Island life.

Webb, C. O. 2000. Exploring the Phylogenetic Structure of Ecological Communities: An Example for Rain Forest Trees. - Am. Nat. 156: 145–155.

Webb, C. O. et al. 2002. Phylogenies and Community Ecology. - Annu. Rev. Ecol. Syst. 33: 475–505.

Webb, C. O. et al. 2008. Phylocom: software for the analysis of phylogenetic community structure and trait evolution. - Bioinformatics 24: 2098–2100.

Weigelt, P. and Kreft, H. 2013. Quantifying island isolation - insights from global patterns of insular plant species richness. - Ecography 36: 417–429.

Weigelt, P. et al. 2013. Bioclimatic and physical characterization of the world’s islands. - Proc. Natl. Acad. Sci. 110: 15307–15312.

Weigelt, P. et al. 2015. Global patterns and drivers of phylogenetic structure in island floras. - Sci. Rep. in press.

Whittaker, R. J. and Fernández-Palacios, J. M. 2007. Island biogeography: ecology, evolution, and conservation.

Wilson, E. O. 1969. The species equilibrium. - In: Diversity and Stability in Ecological Systems. Systems Brookhaven National Laboratory Upton. Brookhaven Symposia in Biology.

