## Supplementary result 1 for "Tracking the origin of island diversity: insights from divergence with the continental pool in monocots"

**Appendix S1 : Relationship between solar radiation, latitude and phylogenetic divergence**

A) Mean annual solar radiation in function of latitude

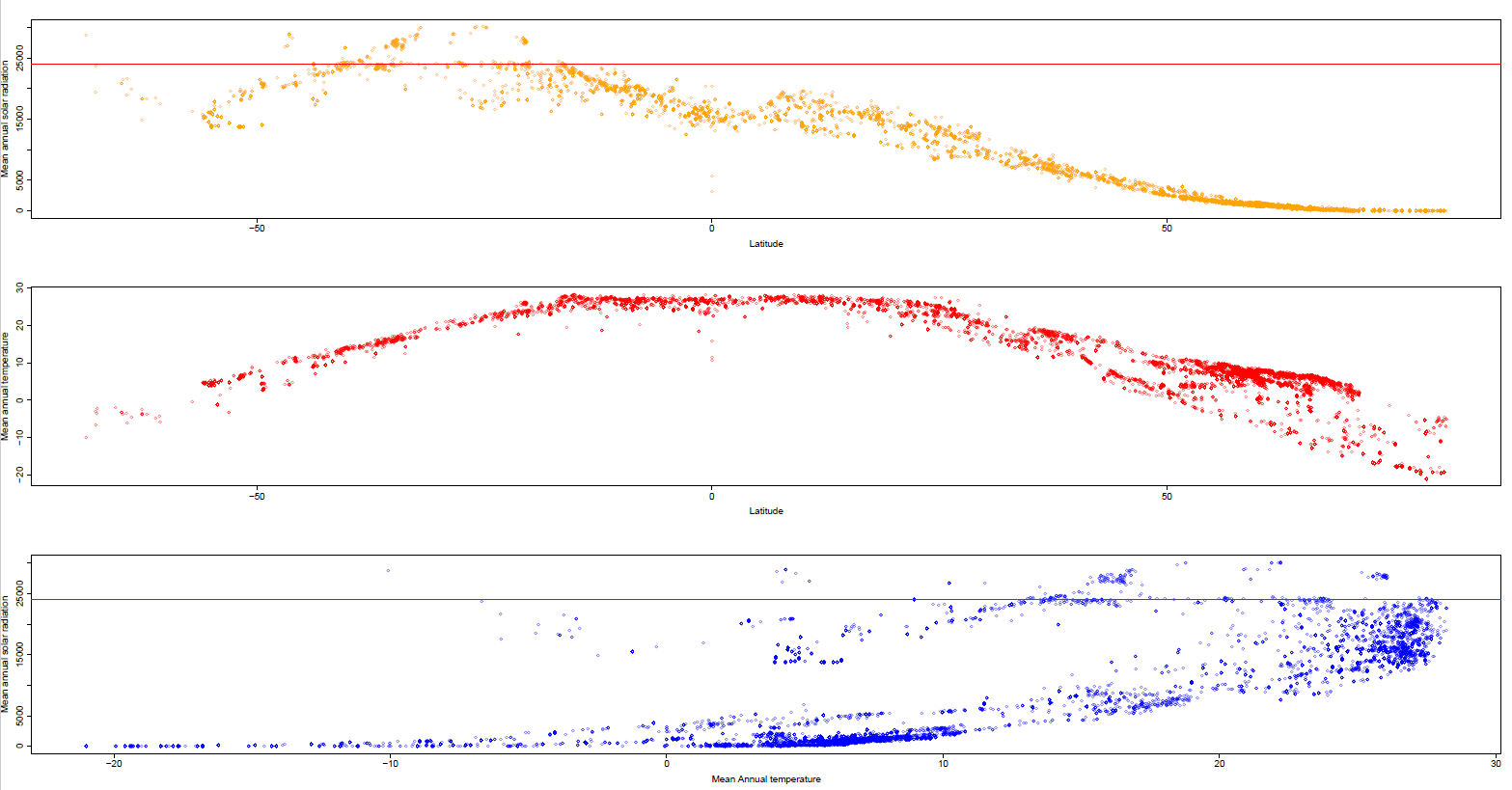

**Figure S1** : a) Mean annual solar radiation in function of latitude b). Mean annual temperature in function of latitude c) Mean annual solar radiation in function of temperature

Graph a) shows two peaks of solar radiation in function of latitude, both centered on the tropics. The red line shows that a similar amount of solar radiation can occur at distinct latitudes. It shows that some areas could diverge because they are located at different latitudes receiving however the same amount of solar radiation. Phylogenetic divergence due to differences in latitude may be due to past and current climate, geologic history, that influenced diversification. Graph b) displays the variation in temperature in function of latitude. Temperature tend to be homogenous in the tropics. Consequently, some islands and continental areas situated at different latitudes in the tropics, and with high phylogenetic divergence, may have a very low difference in mean annual temperature. Graph c) shows that islands receiving similar amount of solar radiation can have a different mean annual temperature.

B) Effect of difference in latitude and longitude between islands and continents on phylogenetic divergence

1) Oceanic islands

|  | A) ***10 nearest continental areas*** | B) ***5 nearest continental areas*** | ***C) 20 nearest continental areas*** |
| --- | --- | --- | --- |
| *Difference in latitude* ***(MPD_β_)*** | +*** | +*** | +*** |
| *Difference in latitude* ***(MNTD_β_)*** | +*** | +*** | +*** |
| *Difference in longitude* ***(MPD_β_)*** | +*** | +*** | +*** |
| *Difference in longitude* ***(MNTD_β_)*** | +*** | +*** | +*** |

2) Continental islands

|  | A) ***10 nearest continental areas*** | B) ***5 nearest continental areas*** | C) ***20 nearest continental areas*** |
| --- | --- | --- | --- |
| *Difference in latitude* ***(MPD_β_)*** | +*** | +*** | +*** |
| *Difference in latitude* ***(MNTD_β_)*** | +*** | +*** | +*** |
| *Difference in longitude* ***(MPD_β_)*** | +*** | +*** | +*** |
| *Difference in longitude* ***(MNTD_β_)*** | +*** | +*** | +*** |

**Tables S1.** Effect of the difference in latitude estimated from generalized linear models 1) Oceanic island 2) Continental island

C) Effect of difference in solar radiation and temperature on phylogenetic divergence in tropical and non-tropical islands

1) Oceanic islands

| ***a) MPD_β_*** | ***A) 10 nearest continental areas*** | | ***B) 5 nearest continental areas*** | | ***C) 20 nearest continental areas*** | |
| --- | --- | --- | --- | --- | --- | --- |
|  | Non-tropical | Tropical | Non-tropical | Tropical | Non-tropical | Tropical |
| *Solar radiation difference* | - | -*** | + | -*** | -*** | -*** |
| *Temperature difference* | +*** | -*** | - | -*** | +*** | -*** |
| ***b) MNTD_β_*** |  |  |  |  |  |  |
| *Solar radiation difference* | +*** | -*** | +*** | -** | +*** | - |
| *Temperature difference* | +*** | -. | +*** | - | +*** | -*** |

2) Continental islands

| ***a) MPD_β_*** | ***A) 10 nearest continental areas*** | | ***B) 5 nearest continental areas*** | | ***C) 20 nearest continental areas*** | |
| --- | --- | --- | --- | --- | --- | --- |
|  | Non-tropical | Tropical | Non-tropical | Tropical | Non-tropical | Tropical |
| *Solar radiation difference* | -*** | -*** | +*** | +** | -*** | +* |
| *Temperature difference* | +*** | -*** | +** | - | +*** | -*** |
| ***b) MNTD_β_*** |  |  |  |  |  |  |
| *Solar radiation difference* | +*** | - | +*** | +*** | +*** | -*** |
| *Temperature difference* | +*** | -*** | +** | +** | +*** | +*** |

**Tables S3.** Effect of difference in solar radiation and temperature estimated from datasets where tropical islands were excluded (i.e. islands whose latitude is between -30° and 30°) and where only tropical islands were included. 1) Oceanic island 2) Continental island

**Conclusion**

Effect of environment on divergence may be related to the geographic position of islands and especially latitude. This may be illustrated by the unexpected finding that in areas with high difference in solar radiation deep branches co-occur more frequently than in areas with similar insolation conditions. This was surprising because differences in solar radiation may act as a strong climatic barrier due to differences in energy supply between areas so that only evolutionary distant lineages are expected to co-occur on islands and continents. Our result is thus counter-intuitive and a possible explanation maybe found in the relationship between solar radiation and latitude. Actually, solar radiations show a latitudinal gradient centered on the tropics and not on the equator. This relationship between latitude and solar radiation is probably due to the angle of solar radiation that becomes more oblique at high latitudes (25). From this, mainly in the tropics, some islands and continents situated at very different latitudes are similarly exposed to solar radiations. Difference in latitude implies difference in species and lineage composition for a number of reasons, including current and past climate, specialization, low dispersal but also evolutionary diversification, which have been explored elsewhere and are still investigated (25). A similar effect of difference in latitude can be evidenced in the occurrence of high divergence between areas with low temperature difference but situated at different latitudes in the tropics.
