## Supplementary result 2 for "Tracking the origin of island diversity: insights from divergence with the continental pool in monocots"

|  |  | **MPDα** | **meanED** | **sdED** | **Nb top 10 ED** | **Proportion of top10 ED species** | **Genus richness** | **VPD** |
| --- | --- | --- | --- | --- | --- | --- | --- | --- |
| Geographic variables | Area |  |  |  |  |  |  |  |
|  | SLMP |  |  |  |  |  |  |  |
|  | Elevation |  |  |  |  |  |  |  |
|  | Min.Dist |  |  |  |  |  |  |  |
|  | Latitude |  |  |  |  |  |  |  |
|  | Longitude |  |  |  |  |  |  |  |
| Climatic variables | Number of ecosystems |  |  |  |  |  |  |  |
|  | Temperature |  |  |  |  |  |  |  |
|  | Rainfall |  |  |  |  |  |  |  |
|  | Rainfall seasonality |  |  |  |  |  |  |  |
|  | Wind speed |  |  |  |  |  |  |  |
|  | Sd. In vapour pressure |  |  |  |  |  |  |  |
| Historical variables | Velocity of past climate change |  |  |  |  |  |  |  |
|  | GMMC |  |  |  |  |  |  |  |
| Sampling effort | ICE |  |  |  |  |  |  |  |

**Appendix S2 : Effect of island features on phylogenetic structure**

**Table S4** : Significance of the relationship between island features and measures of island phylogenetic structure was estimated thanks to generalized linear models. Variable responses were measures of phylogenetic structure taken independently and explanatory variables were island features. These resulted in 7 models corresponding to the number of phylogenetic structure measures.

[expli](http://www.wsl.ch/staff/niklaus.zimmermann/papers/PPEES_Smycka_2016.pdf)

Slight significant negative effect p-val<0.01

High significant negative effect p-val<0.001

Slight significant positive effect p-val<0.01

High significant positive effect p-val<0.001
