## Supplementary result 3 for "Tracking the origin of island diversity: insights from divergence with the continental pool in monocots"

### BRT contributions of spatial and environmental distance to phylogenetic divergence

#### a) Oceanic islands

##### MPD$_\beta$

*Dist= Spatial distance*
*Dist_env= Euclidean environmental distance*
*Diff_temp=Difference in mean annual temperature*
*Diff_rainf= Difference in mean annual rainfall*
*Diff_srad= Difference in mean annual solar radiation*
*Diff_wind=Difference in mean annual wind speed*
*Nb.diff.ecoreg=Number of different ecoregions*

*10 closest continental areas*

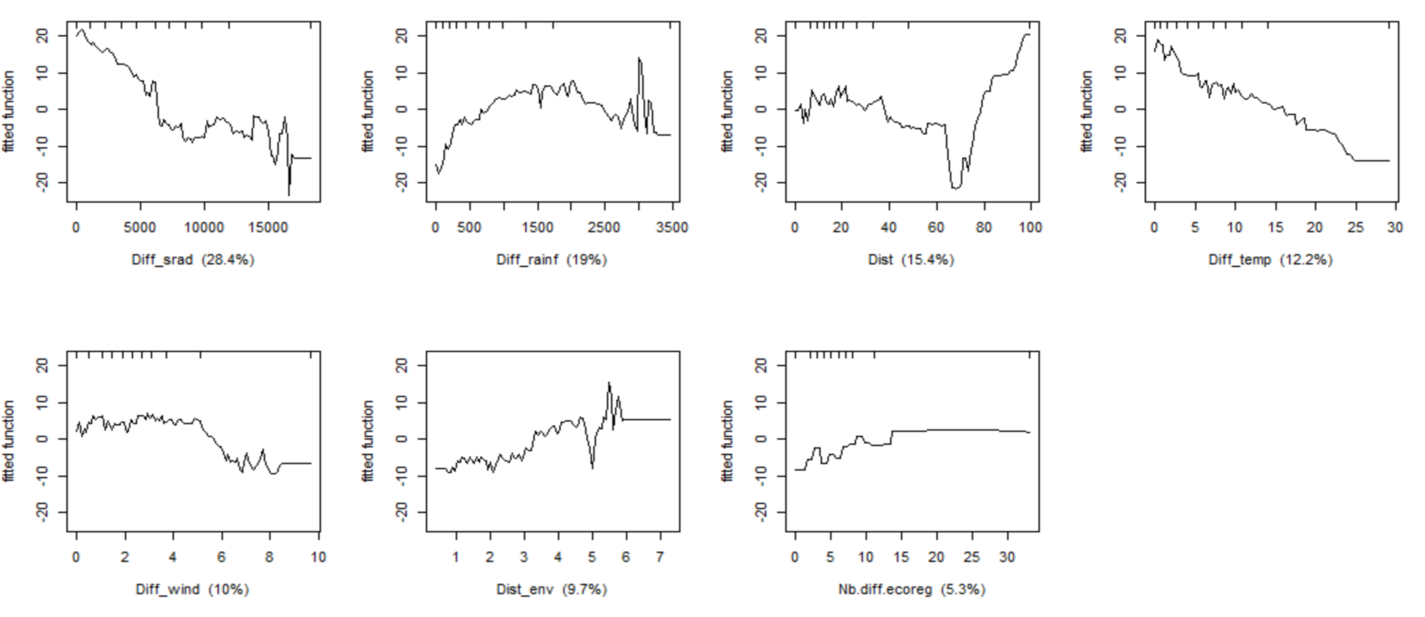

*5 closest continental areas*

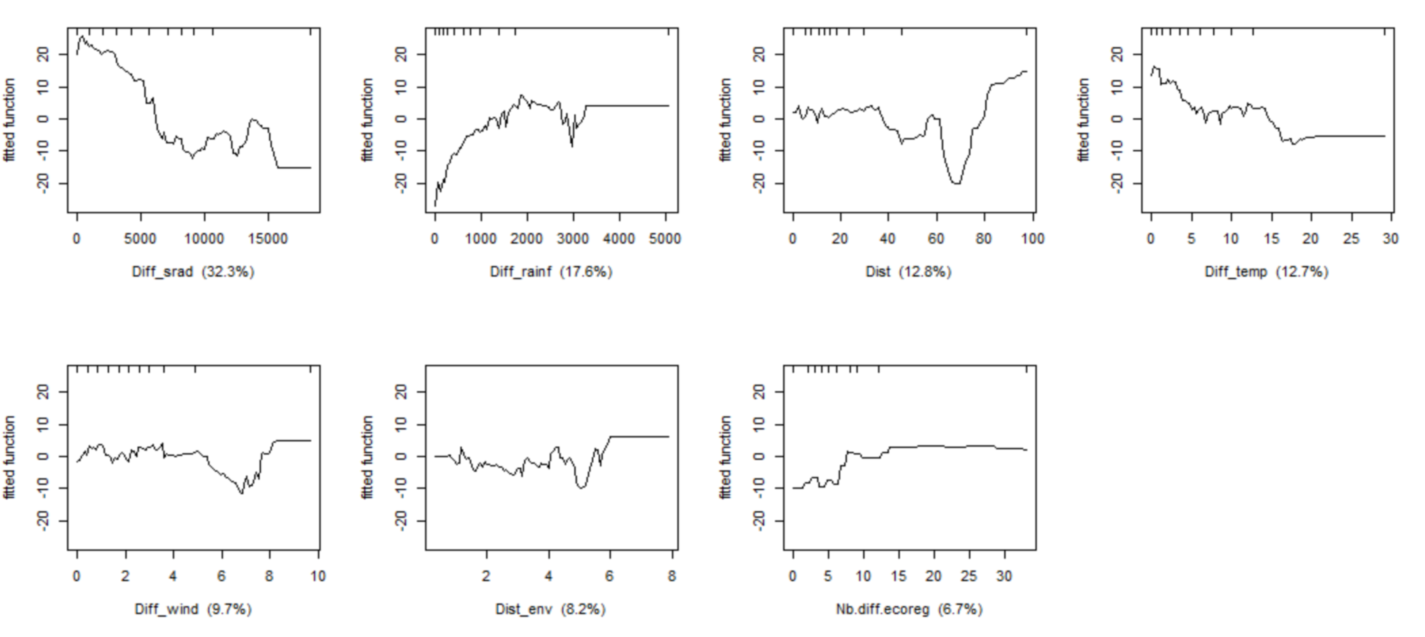

*20 closest continental areas*

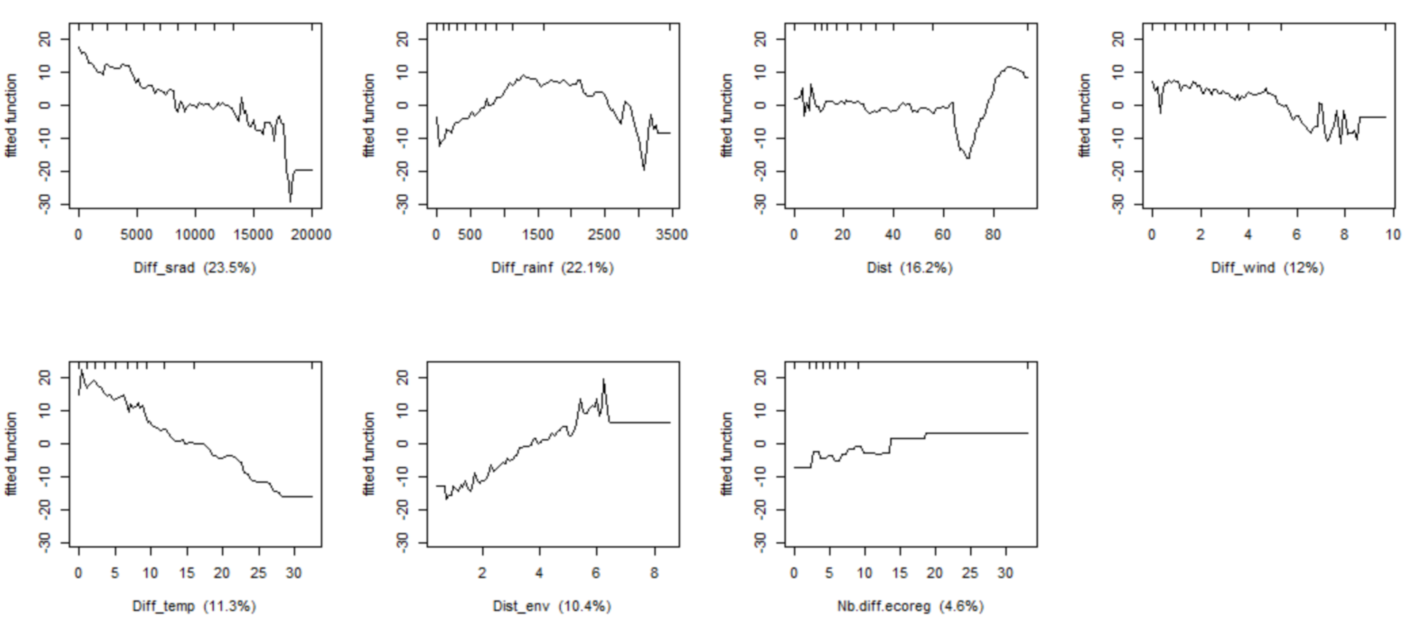

##### MNTD$_\beta$

*10 closest continental areas*

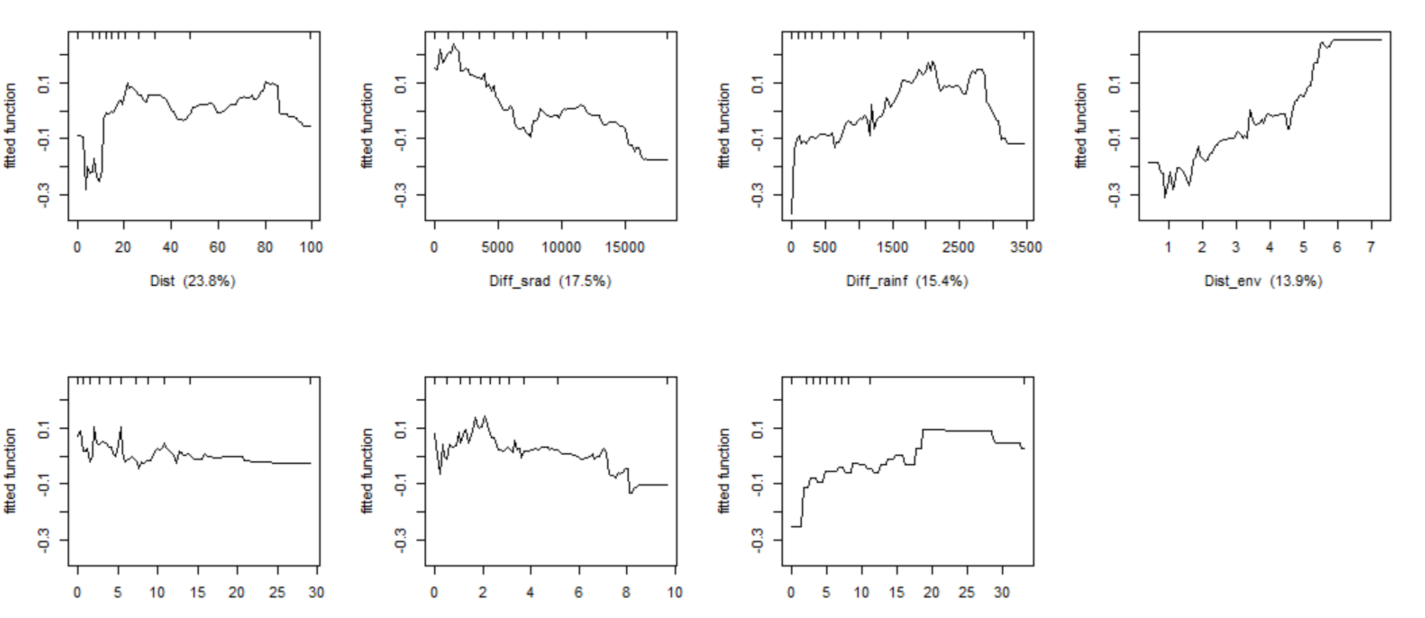

*5 closest continental areas*

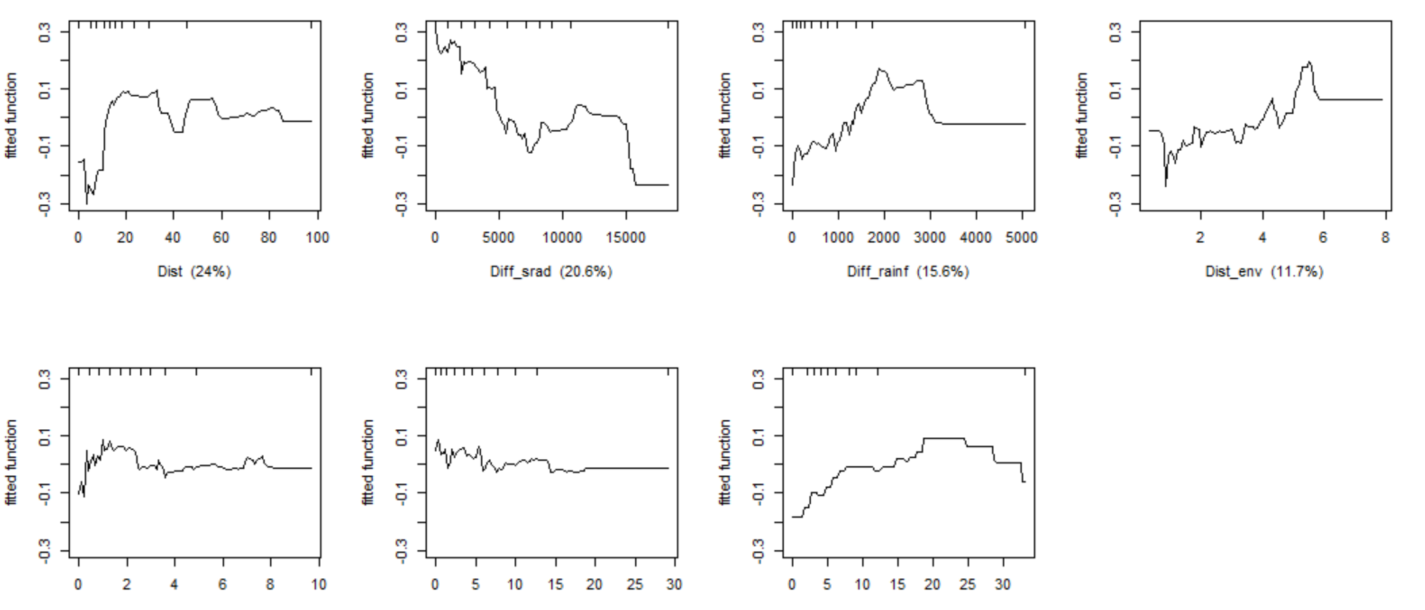

*20 closest continental areas*

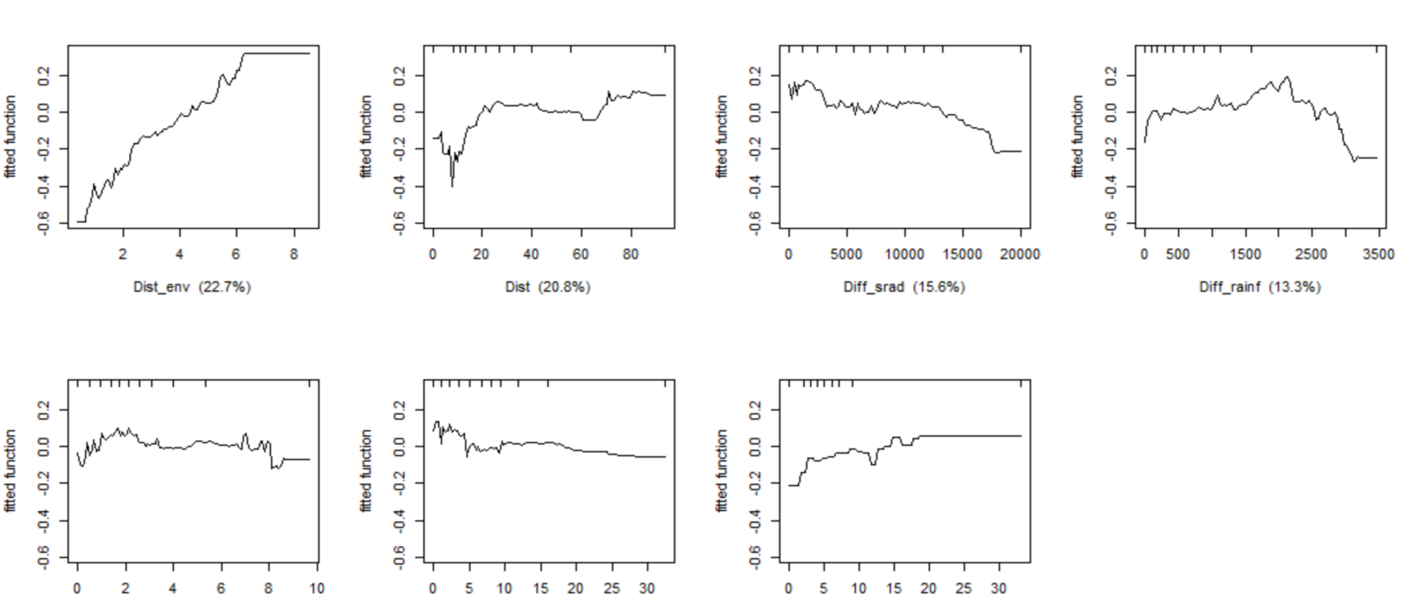

#### b) Continental islands

##### MPD$_\beta$

*10 closest continental areas*

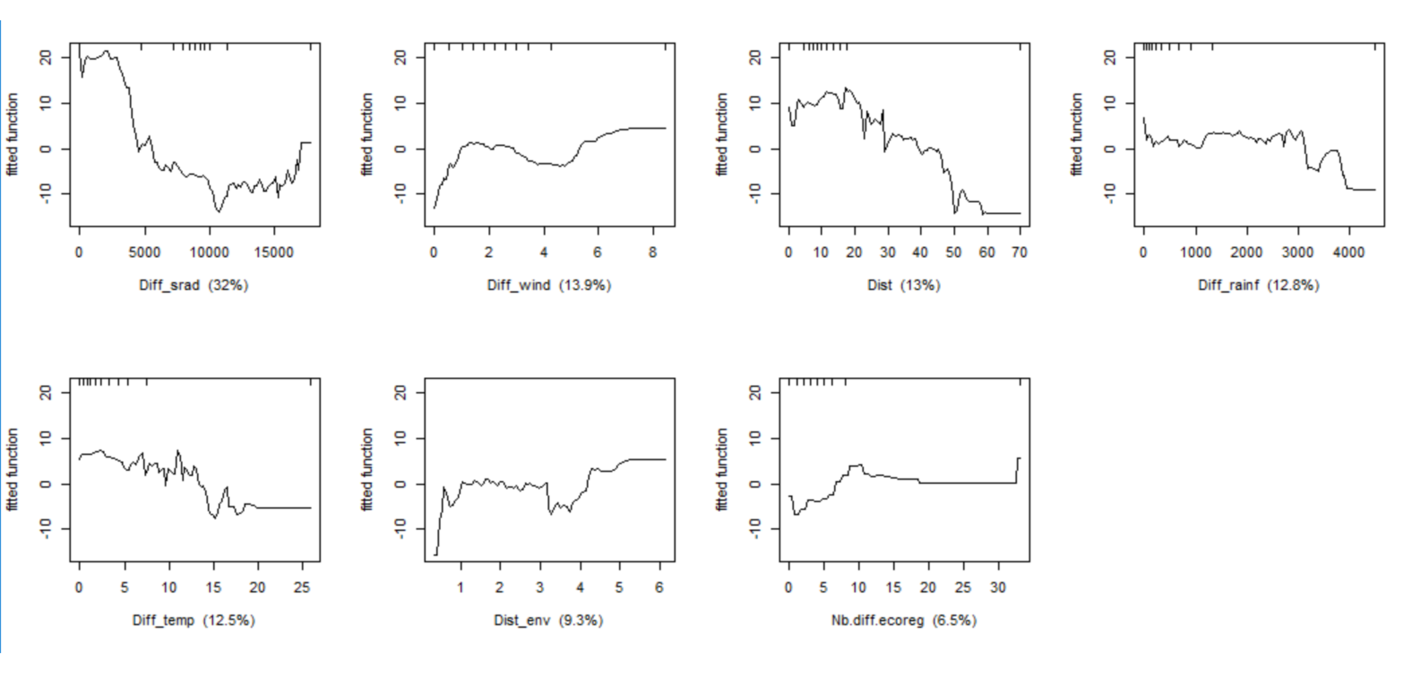

*5 closest continental areas*

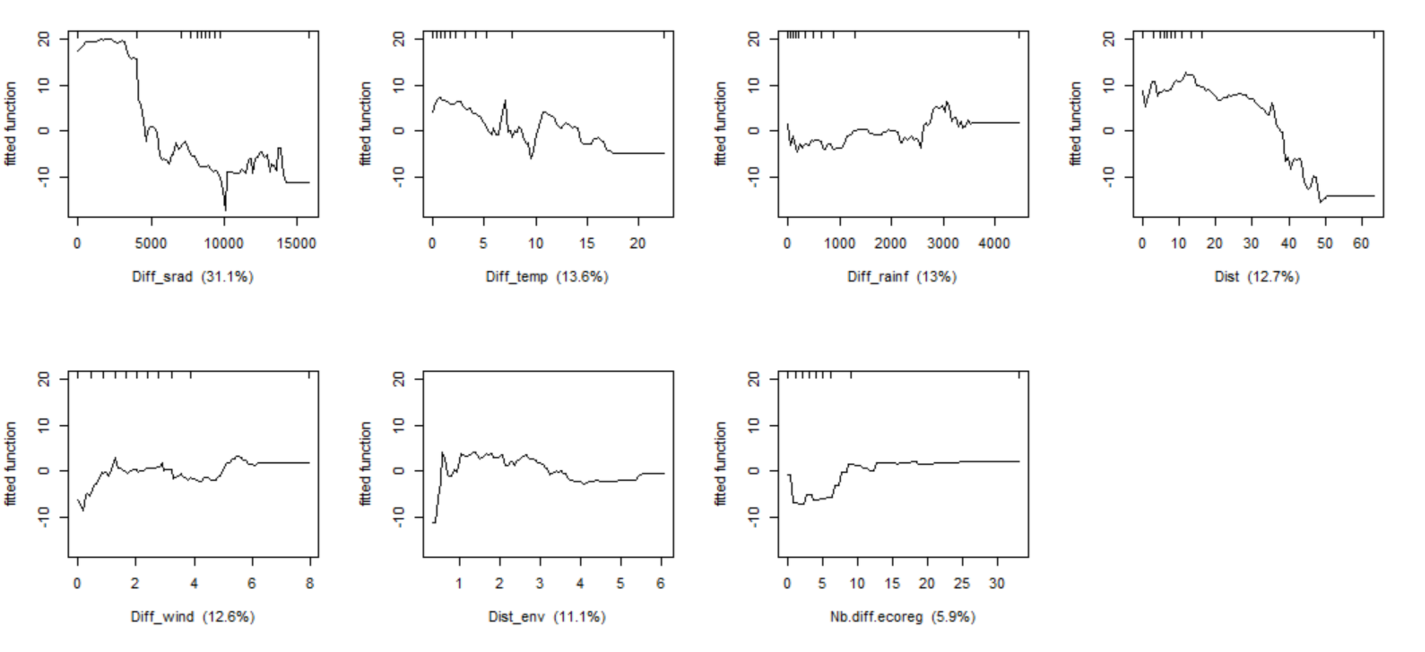

*20 closest continental areas*

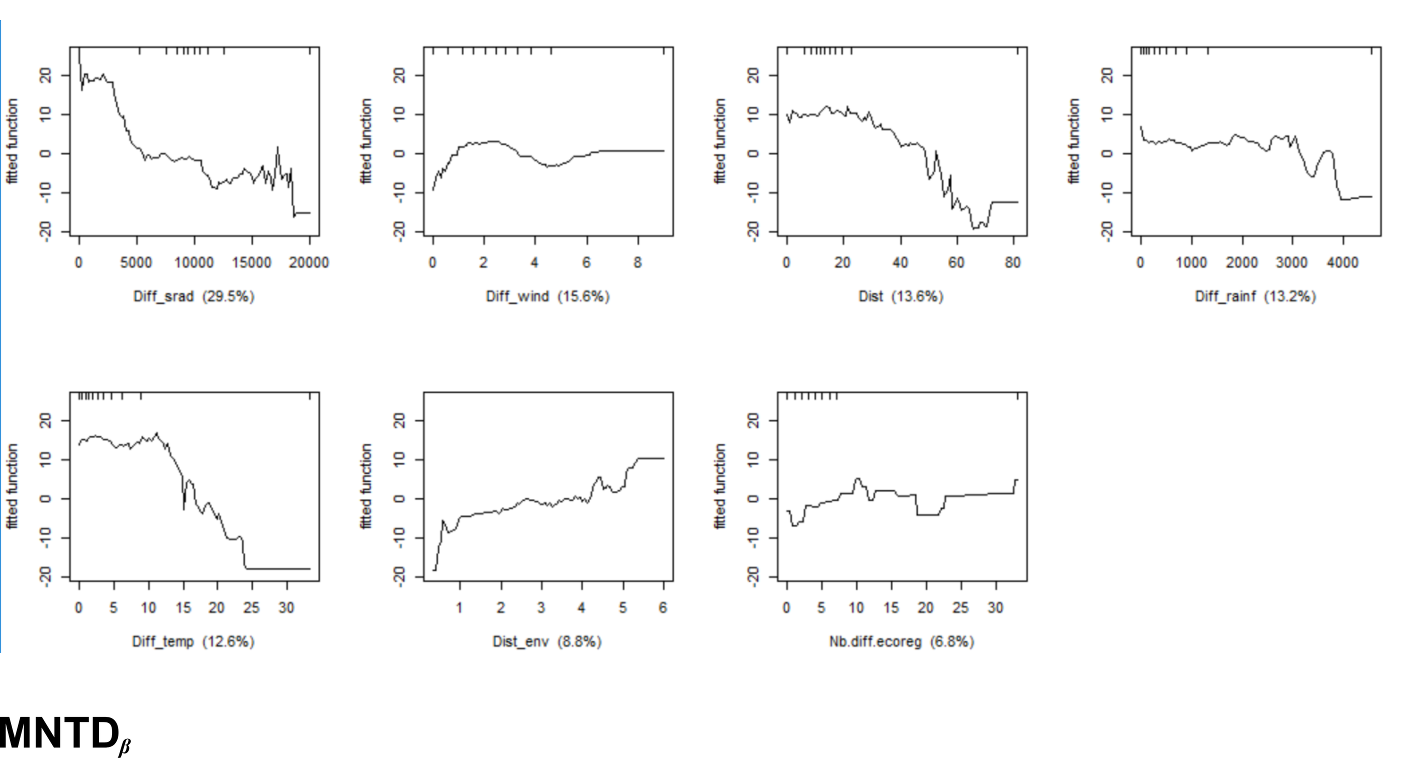

##### MNTD$_\beta$

*10 closest continental areas*

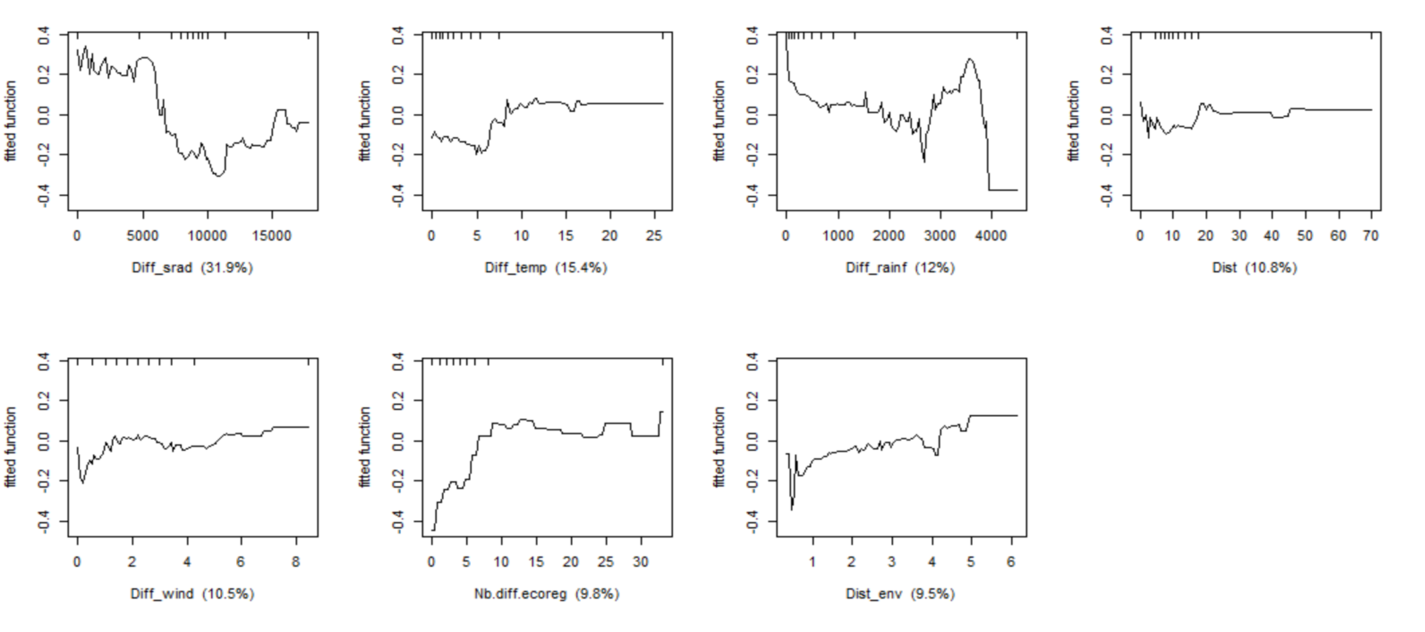

*5 closest continental areas*

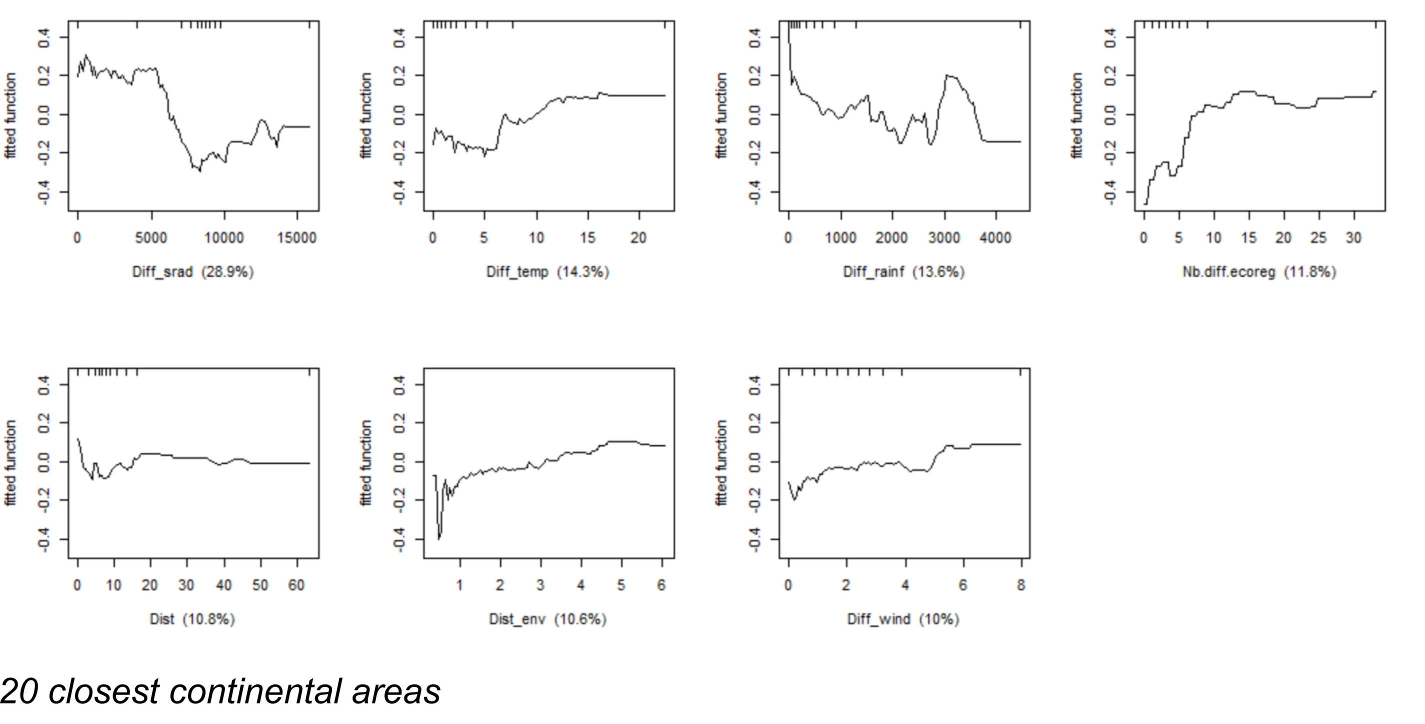

*20 closest continental areas*

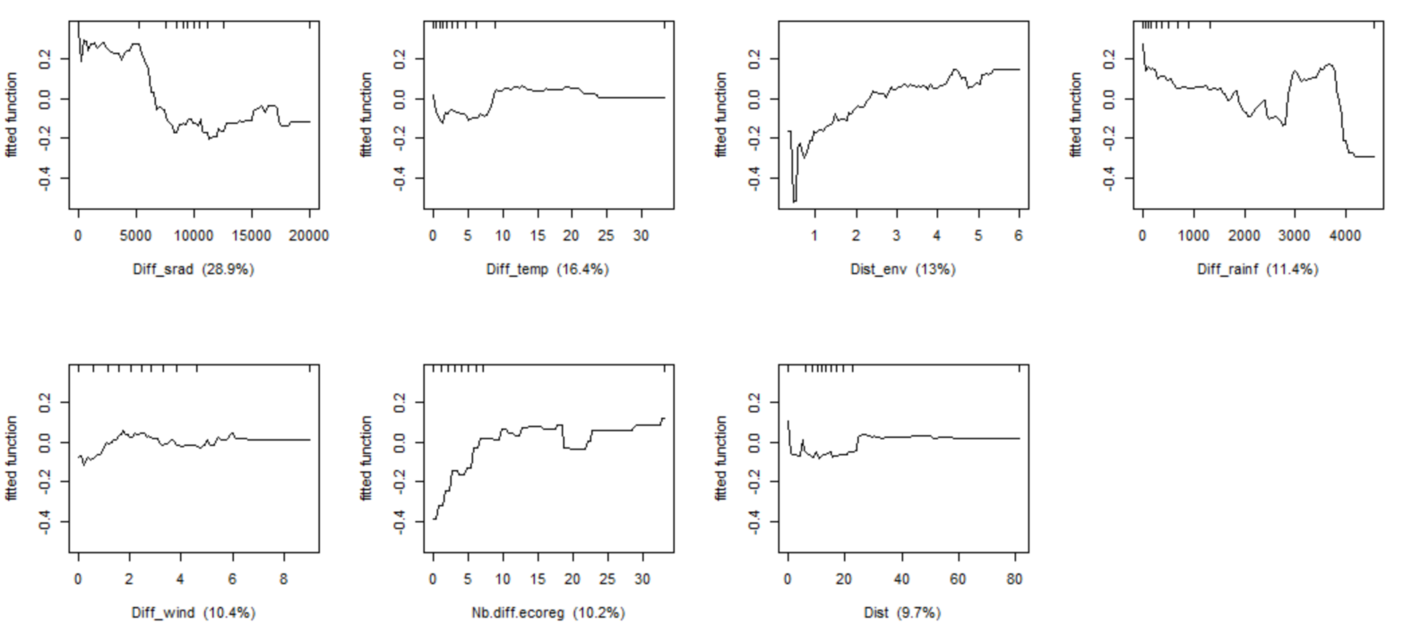
